# Functional subtypes of rodent melanopsin ganglion cells switch roles between night and day illumination

**DOI:** 10.1101/2023.08.26.554902

**Authors:** Michael H. Berry, Joseph Leffler, Charles N. Allen, Benjamin Sivyer

## Abstract

Intrinsically photosensitive retinal ganglion cells (ipRGCs), contain the photopigment melanopsin, and influence both image and non-image forming behaviors. Despite being categorized into multiple types (M1-M6), physiological variability within these types suggests our current understanding of ipRGCs is incomplete. We used multi-electrode array (MEA) recordings and unbiased cluster analysis under synaptic blockade to identify 8 functional clusters of ipRGCs, each with distinct photosensitivity and response timing. We used Cre mice to drive the expression of channelrhodopsin in SON-ipRGCs, enabling the localization of distinct ipRGCs in the dorsal retina. Additionally, we conducted a retrospective unbiased cluster analysis of ipRGC photoresponses to light stimuli across scotopic, mesopic, and photopic intensities, aimed at activating both rod and cone inputs to ipRGCs. Our results revealed shared and distinct synaptic inputs to the identified functional clusters, demonstrating that ipRGCs encode visual information with high fidelity at low light intensities, but poorly at photopic light intensities, when melanopsin activation is highest. Collectively, our findings support a framework with at least 8 functional subtypes of ipRGCs, each encoding luminance with distinct spike outputs, highlighting the inherent functional diversity and complexity of ipRGCs and suggesting a reevaluation of their contributions to retinal function and visual perception under varying light conditions.

## Introduction

Intrinsically photosensitive retinal ganglion cells (ipRGCs)^1,2^, contain the photopigment melanopsin^2,3^, and extend projections to both image-forming and non-image forming regions within the brain^1,2,4–9^, impacting a diverse array of behaviors^8,10–17^. The axons of ipRGCs form the principle retinal projection to the suprachiasmatic nucleus, the brain’s master circadian pacemaker, where they control light-mediated entrainment of circadian rhythms. ipRGCs also project to the olivary pretectal nucleus where they influence the pupillary light reflex^2,18^. In addition to non-image forming visual areas, ipRGCs project to brain regions such as the lateral geniculate nucleus influencing image-forming regions. Presently, ipRGCs are categorized into six distinct ‘types’ (M1-M6), primarily defined by their dendritic morphology and melanopsin expression^19,20^. The most extensively researched M1 ipRGCs exhibit simplistic dendritic structures and express the highest concentration of melanopsin, thereby yielding the greatest intrinsic photosensitivity^21^. In contrast, non-M1 ipRGCs (M2-M6)^10,22–25^, possess unique morphologies, including more intricate dendritic structures, and exhibit weaker intrinsic photoresponses. However, they receive more substantial synaptic input from upstream bipolar cells, which transmit rod- and cone-derived signals^26,27^.

While the morphology of M1-M6 ipRGCs has been well-defined, increasing evidence suggests functional variability within these types. M1 ipRGCs can be split based on their expression of the transcription factor Brn3b, leading to different patterns of central innervation and thus separate functional controls over photo-entrainment and the pupillary light reflex^13^. M1 ipRGCs have diverse intrinsic photoresponses^19,28–30^ and appear to fall into discrete groups based on the effect of depolarization block^31^. These variable intrinsic photoresponses and spiking mechanisms allow M1 ipRGCs to encode luminance across a broad range of light intensities^29,32^. M1 ipRGCs also differ based on their rod-mediated photoreceptor drive, with a gradient of rod-mediated input to ipRGCs that scales with their dendritic complexity and overlays with variable intrinsic photosensitivity^28^. Recent results suggest that M1 ipRGCs differ in their neurotransmitter release^33^, and location within the retina, with some forming tiling mosaics^9,34^. This suggests they likely fall into discrete functional subtypes, although it remains unclear how their functional diversity is represented at a population level.

The transcriptional heterogeneity of M1 ipRGCs implies they likely divide into at least two unique clusters, known as M1a and M1b^35^. These classifications could align with ipRGCs receiving varied synaptic inputs and distinct light response features^26,28,29^. Further clusters corresponding to M2 and M4 ipRGCs are apparent when using the expression of OPN4 as a transcriptional marker to identify ipRGCs. However, this method is not effective in differentiating M3, M5, and M6 ipRGCs, some of which may belong to a single cluster (Cluster C22). Another investigation into the transcriptional diversity of ipRGCs reveals that many ipRGC types might express specific genes, potentially affecting their physiological light responses^36^. The correlation between these varied transcriptional profiles and the functional types of ipRGCs within the broader population remains undefined.

Despite less knowledge about the functional heterogeneity of non-M1 ipRGCs, recent findings suggest the existence of similar variation. ipRGCs demonstrate a variety of intrinsic light responses due to differences in melanopsin expression and variations in their second messenger systems^37–40^. Compared to M1 ipRGCs, M2 and M4 ipRGCs employ diverse sets of second messenger cascades subsequent to melanopsin, leading to the distinct activation of multiple effector channels^8,37,41^. This difference likely constitutes another mechanism contributing to the diversity of intrinsic photoresponses within ipRGC types. Nonetheless, how this functional diversity of ipRGCs presents itself on a population level remains uncertain. Furthermore, it remains unclear how the functional diversity among all ipRGCs is driven by their intrinsic photoresonses, their differing synaptic inputs, or a mixture of both.

We aimed to classify the functional diversity of ipRGCs in the retina using unique techniques to identify them in isolated preparations of mouse retina. We hypothesize that unique functional subgroups of ipRGCs can be identified based on their photoresponses both driven by conventional photoreceptor inputs and by intrinsic melanopsin signaling. We utilized multi-electrode array (MEA) recordings and unbiased cluster analysis under synaptic blockade to identify eight functional clusters of ipRGCs, each with distinct photosensitivity and response timing. By developing a technique to optotag channelrhodopsin2(ChR2)-expressing ipRGCs, we were able to localize photoresponses from unique ipRGCs subtypes. We conducted a retrospective analysis of ipRGC photoresponses prior to synaptic blockade at various light intensities, encompassing scotopic, mesopic, and photopic ranges. This range of visual stimuli activated both rod and cone inputs to ipRGCs, thereby revealing the nature of both shared and distinct synaptic inputs to the functional clusters identified. Our results suggest ipRGC diversity is driven primarily by differences in their intrinsic photosensitivity and not their photoreceptor-mediated synaptic inputs.

We also examined the encoding of visual information across these differing light-intensities. Many ipRGCs project to the LGN and therefore influence image-forming vision. In M4 ipRGCs, melanopsin contributes to the visual responses at low light intensities encompassing scotopic illumination and increases the contrast sensitivity of these ipRGCs and their associated behaviors^8,10^. However, the encoding of visual information by all ipRGCs across scotopic to photopic illumination in the absence of synaptic blockade remains unclear. Our results illustrate most ipRGCs encode visual information with high fidelity at low light intensities, but poorly at photopic light intensities, where melanopsin activation is at its peak.

This pattern of responses hints at the functional adaptability in ipRGCs, suggesting that they may switch roles depending on environmental luminance. These insights not only underscore the complexity and functional diversity of ipRGCs but also prompt a reevaluation of their contributions to retinal function and visual perception under varying light conditions.

## Results

### MEA recordings from ipRGCs populations in the mouse retina

Despite the fact that most studies on the functional diversity of ipRGCs have relied on traditional single cell recording techniques, these methods pose significant challenges. ipRGC sensitivity to illumination history^31^, and their activation by both single and two-photon fluorescent light excitation make comparisons of ipRGC photoresponses in single preparations difficult. Even brief and dim illumination can stimulate melanopsin^28,42^ and induce retinal adaptation, thereby altering their physiological properties. To address these sensitivity issues, it became essential for us to characterize ipRGC photoresponses at a population level, by recording light responses using MEA techniques^20,26,27,40,43–49^ (Figure 1; supplementary Fig 1). This allowed us to capture light responses from thousands of RGCs simultaneously, from scotopic to photopic light intensities prior to and during synaptic blockade.

**Figure 1:**
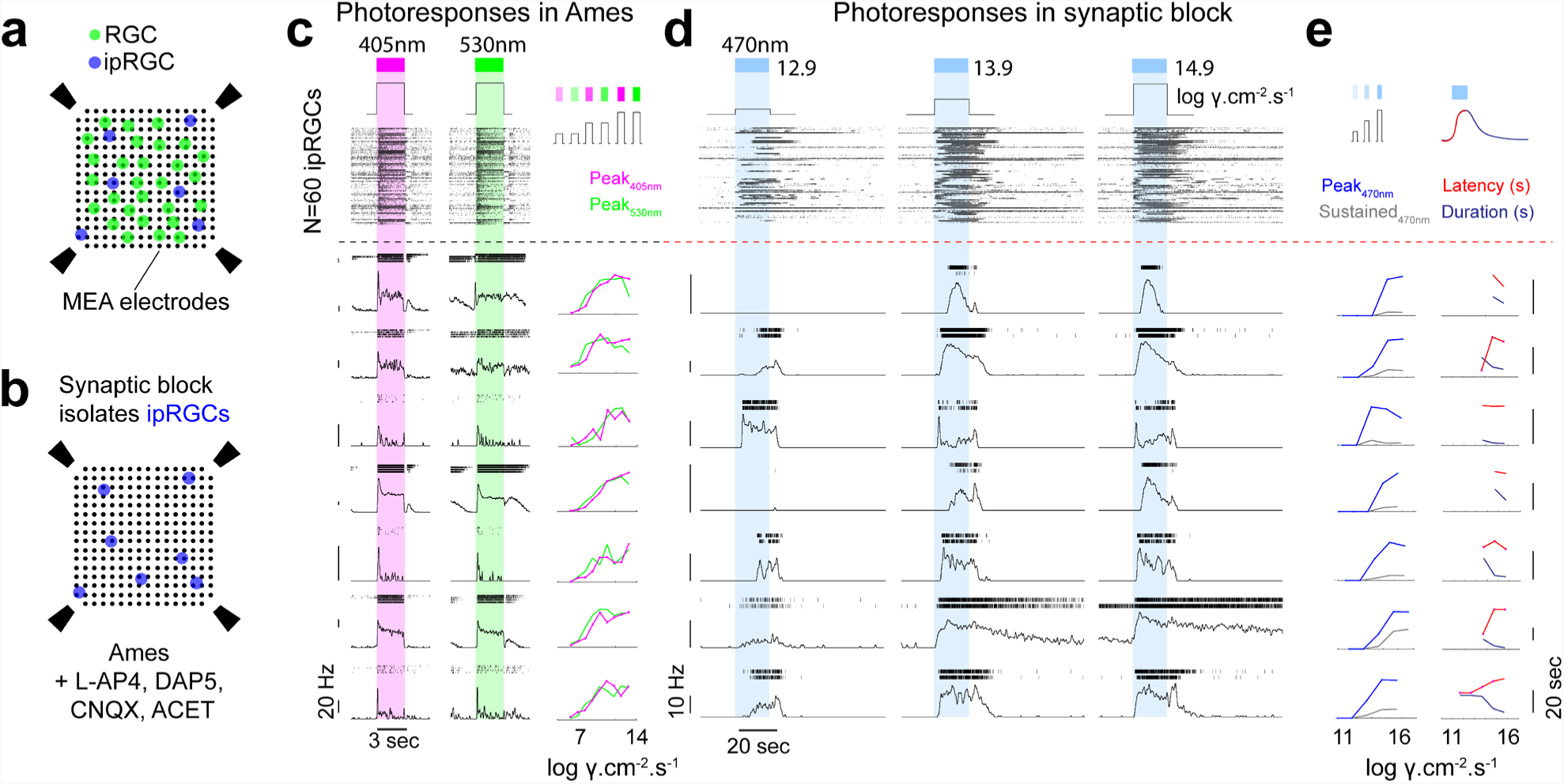
MEA recording configuration and ipRGC response diversity in the mouse retina. **(a)** A 256 channel MEA records spike responses from both RGCs and ipRGCs. **(b)** Synaptic block using L-AP4, D-AP5, CNQX and ACET blocks photoreceptor-mediated synaptic transmission and isolates light responses from ipRGCs. **(c)** Representative examples of dark-adapted photoresponses in 7 ipRGCs in Ames and, **(d)** in the presence of synaptic blockers. Individual cells exhibit variability in rod and cone-driven responses to UV (left) and green (right) light (c) and variability in intrinsic sensitivity (left and middle), latency, and duration of responses (right) to blue light under synaptic blockers.

Under dark-adapted conditions, we recorded photoresponses to full-field UV (405nm) and green (530nm) light for all RGCs (Figure 1a,c), including chirp stimuli to probe temporal and contrast-dependent responses^50^. Following these initial recordings, we added synaptic blockers to the solution and identified ipRGCs by their inherent photosensitivity to blue (470nm) light (Figure 1b,d).

Individual units, each representing a single ipRGC, showed substantial variability in their light-induced spiking in response to 20-second incremental light stimuli (13-15 log photons cm-2 s-1; Figure 1d). This included different photosensitivity thresholds, peak or sustained firing rates (Figure 1e) and variable latency and duration of their light responses (Figure 1e). These observations illustrate the diversity of intrinsic photoresponses amongst identified ipRGCs.

To selectively target unique ipRGC populations, including SON-ipRGCs^9^ and other non-M1 ipRGCs, we used *GlyT2^Cre^;Ai32* mice which promotes ChR2 expression in dorsal ipRGCs, SON-ipRGCs and Cre-expressing ipRGCs which are primarily M2 ipRGCs^9^ (Figure 2 a,b). MEA recordings in *GlyT2^Cre^;Ai32* mice were restricted to the dorsal retina and we assumed that our recordings sampled spike responses from most ipRGC types in this region. We hypothesized that we can isolate M1 and M2 ipRGCs based on their stronger intrinsic photoresponses whereas other ipRGC functional groups should have distinctly weaker intrinsic photoresponses^19^.

**Figure 2:**
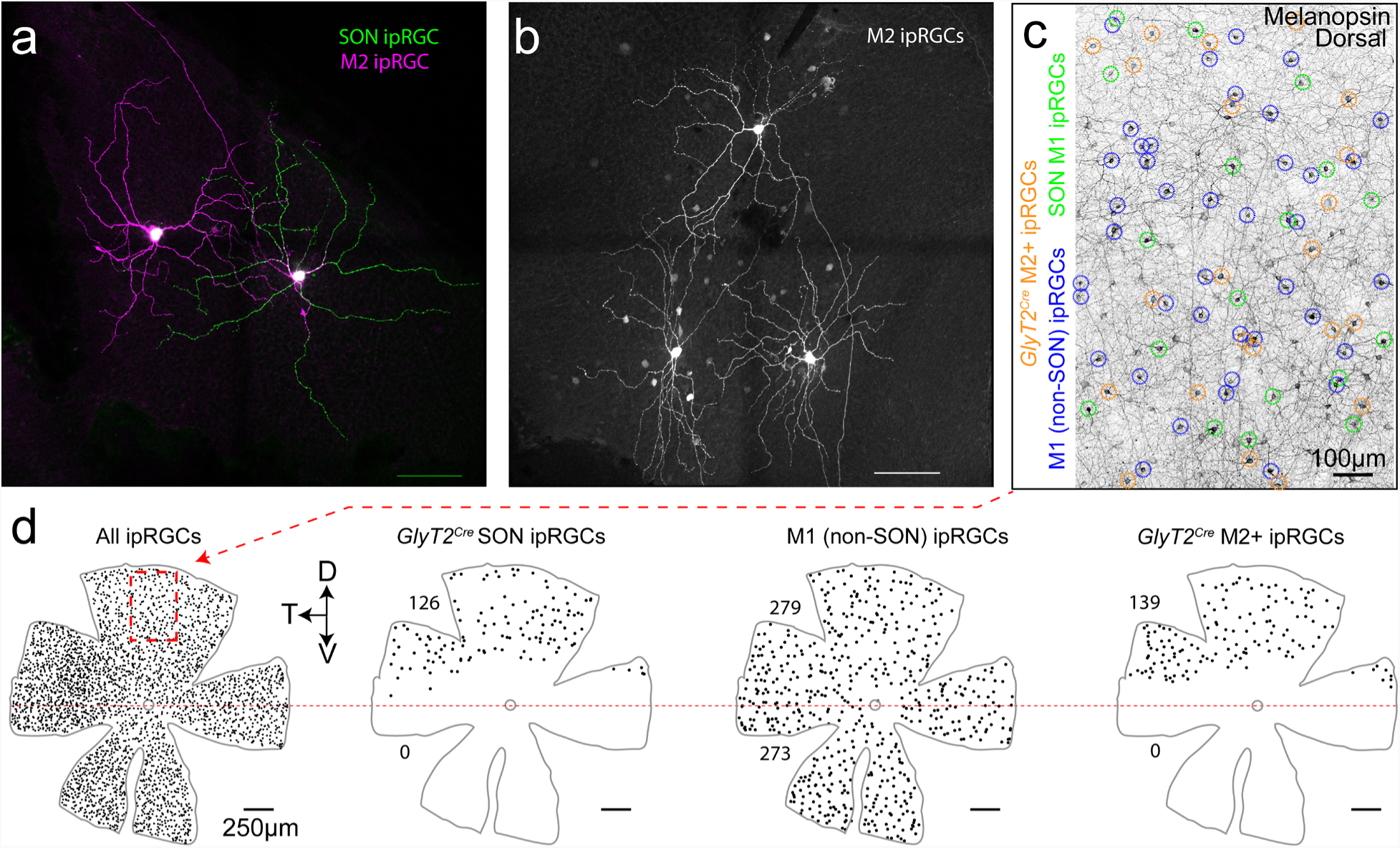
Primary ipRGC subtypes in *GlyT2^Cre^* retinas. **(a)** Confocal micrograph of Neurobiotin-filled SON ipRGC and M2 ipRGC with their dendrites pseudocolored based on their stratification in sublamina-a (green) or sublamina-b (magenta) of the inner plexiform layer. These ipRGCs were targeted in *GlyT2^Cre^;Ai9* mice using epifluorescence. **(b)** Three surrounding Neurobiotin-filled M2 ipRGCs targeted in *GlyT2^Cre^;Ai9* mouse retina illustrate they form a mosaic. **(c)** Confocal micrograph of melanopsin-labeled retina with SON M1 (green), GlyT2*Cre* M2+ (orange) and M1 (non-SON) ipRGCs (blue) identified by their dendritic lamination using melanopsin antibody staining **(d)** Maps of ipRGCs in *GlyT2^Cre^;Ai9* retina illustrating the distribution of all ipRGCs, SON-ipRGCs, M1 ipRGCs and M2+ ipRGCs in the dorsal retina.

To target SON-ipRGCs while also recording from all ipRGCs as a population, we combined MEA recordings with an optotagging strategy. Using *GlyT2^Cre^;Ai32* mice we restricted ChR2-eYFP expression to the SON-ipRGCs in the dorsal retina (Figure 3a). First, we recorded light responses to a wide range of visual stimuli in dark-adapted retinas (Figure 3b,e-g; Step 1). Second, a cocktail of synaptic blockers (40μM L-AP4, 50μM DAP5, 50μM CNQX, 2μM ACET) pharmacologically isolated ipRGCs and we recorded intrinsic photoresponses with 20 sec light stimuli at increasing intensities (Figure 3c,h-j; Step 2). Finally, we partially blocked melanopsin responses with an opsinomide^20,51^ and presented high-frequency and high intensity blue light stimuli (18Hz; 5ms pulse duration; 17 log photons/cm^2^/s) to activate ChR2-expressing ipRGCs (Figure 3d,k-o; Step 3), identifying them amongst other ipRGCs.

**Figure 3:**
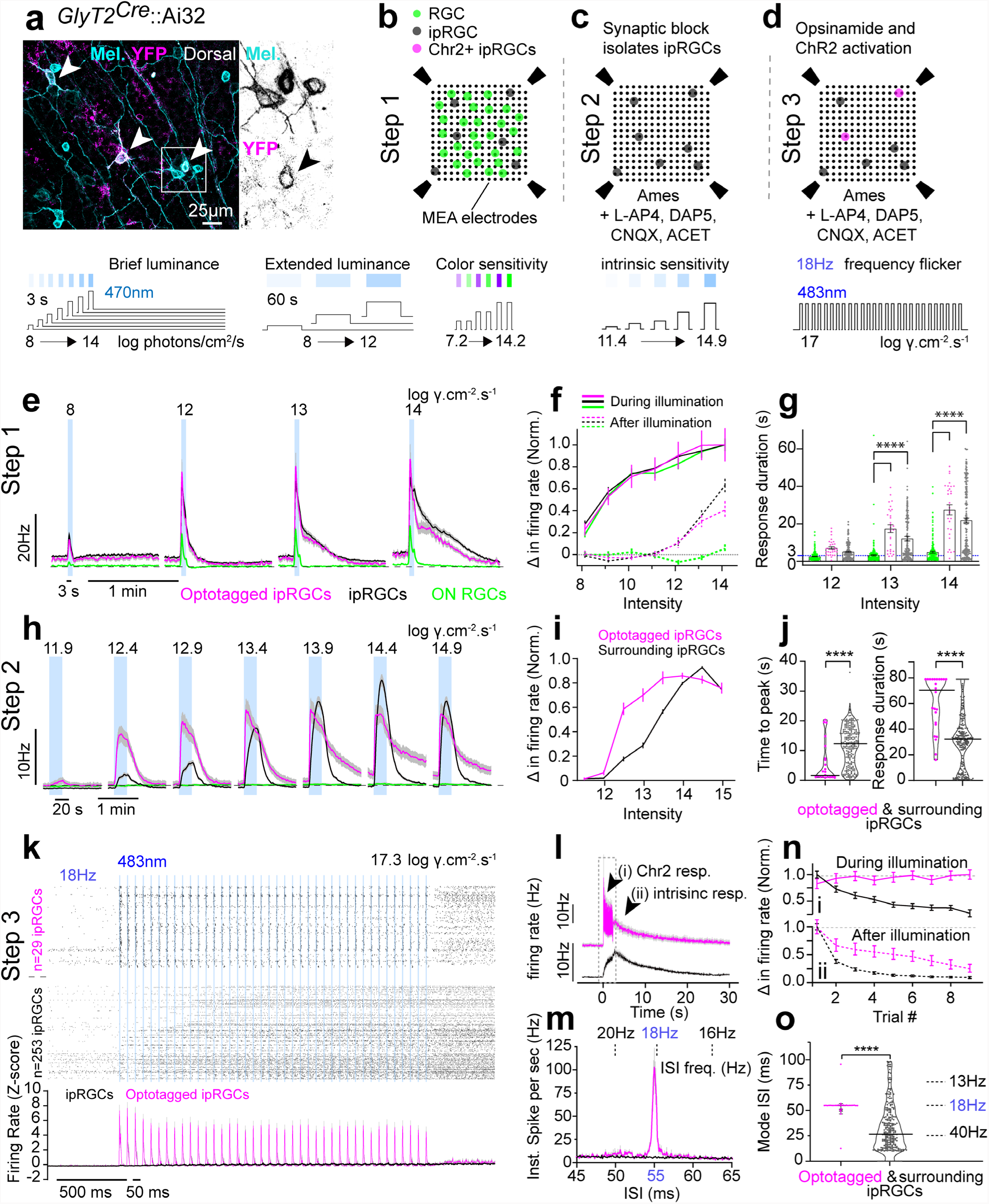
Optotagging of ChR2-expressing ipRGCs. **(a)** Confocal image of GlyT2 ipRGCs co-expressing (white arrows) melanopsin (cyan) and ChR2 (magenta) in *the GlyT2^Cre^;Ai32* dorsal retina. **(b)** Illustration of strategy for recording dark-adapted light responses from ipRGCs on the MEA. Photoreceptor responses are recorded blindly across all RGCs (Step 1). **(c)** intrinsic sensitivity under synaptic blockers identifies ipRGCs (Step 2), and **(d)** high frequency flicker under synaptic blockade and a melanopsin inhibitor (optotagging) identifies ipRGCs that express ChR2 (Step 3). Further analysis allows dark-adapted photoresponses from localized cells (units) to be retroactively examined, see methods for expanded description. **(e-f)** Dark-adapted photoresponse of optotagged ipRGCs (magenta), surrounding ipRGCs (black), and ON RGCs (green) to brief illuminations (3 sec) of blue (470nm) light. Averaged responses (e), normalized change in firing rate (f) and duration of light responses following brief illumination (bar graphs depict mean) at scotopic, mesopic and photopic light intensities. Sustained spiking (beyond the illumination period (e, f - dotted, g – blue dotted) in ipRGCs (magenta & black), compared to ON RGCs (green), emphasizes the role of melanopsin in ipRGC photoresponses. **(h-j)** Intrinsic photoresponses (under synaptic block) from 20 sec blue (470nm) illuminations in optotagged ipRGCs (magenta) and surrounding ipRGCs (black). **(h)** Averaged responses and **(i)** normalized change in firing rate at mesopic to photopic light intensities. **(j)** Violin plots (line depicts median) of time from illumination onset to peak photoresponse (left) and duration of photoresponse (right), measured at 13.9 log photons cm^−2^ s^−1^, highlights the difference in optotagged ipRGCs (magenta) compared to surrounding ipRGCs (black). **(k-o)** Photoresponses of ipRGCs to optotagging high frequency stimulation (18Hz) at 17.3 log photons cm^−2^ s^−1^ under synaptic block and opsinamide AA92593 (melanopsin inhibitor ^20,52^). **(k)** Peri-stimulus time histogram of ipRGCs with (n=29; magenta; optotagged ipRGCs) and without (n=253; black; surrounding ipRGCs) time locked ChR2 responses (N= 4 retina). Raster (top) displays individual spiking responses; histogram displays average change in firing rate to 18Hz flicker normalized by z-score. **(l)** Averaged response of optotagged (magenta) and surrounding ipRGCs (black) with arrows (i,ii) identifying a transient ChR2 dependent component (i; doted box is k) and slow sustained intrinsic (melanopsin dependent) component. **(m)** Tracking of normalized change in firing rate for ChR2 response (i) and intrinsic response (ii) across repeated trials of flicker stimuli (9 trials). **(n)** Average inter-spike interval (ISI) histogram and **(o)** ISI occurrence (mode) in response to the optotagging stimulation. Inter-spike interval frequency equivalents (m-top, o-right) are provided for ease of comparison with stimulation frequency (18Hz). (e-i & k-m) Values are mean ± SEM, (j,o) violin plot bar is median. Statistical significance was assessed using Mann-Whitney test for comparisons between two groups or one-way Anova with Sidaks correction for multiple comparisons (\*\*\*\**p*≤0.0001). N= 282 ipRGCs, N = 4 retina.

These optotagging stimuli isolated a small population of ipRGCs (13%±4.6% of ipRGCs n=4 retina) that had time locked spike responses (Figure 3k - top) with an inter-spike interval equivalent to the stimulation frequency (18Hz; Figure 3m,o). Observing these photoresponses on a larger time scale, all ipRGCs clearly exhibited a slow sustained response (Figure 3l), which resembled melanopsin activation that is not completely blocked by the application of the opsinomide^52^. However, optotagged cells were separated by an additional fast component paired with high frequency stimulation. (Figure 3l; ChR2 resp.). Upon repeated trials, the light induced change in spiking activity of the slow component declined while the ChR2 component remained intact (Figure 3n). These results are consistent with the resistance of ChR2 to photo-bleaching over repeated trials and melanopsin’s capacity for adaptation under bright illumination^31,53–56^.

Once separated into optotagged and surrounding (non-optotagged) ipRGC populations (Figure 3k-o), light responses to visual stimuli acquired prior to synaptic blockade and optotagging were retroactively analyzed. ipRGCs respond to scotopic, mesopic and photopic light stimuli with fast rod and cone mediated drive over a similar time course as other ON RGCs (Figure 3e-g), yet had sustained spike responses to luminance levels that activated melanopsin (Figure 3e,f). Indeed, ipRGCs often spiked for 30 seconds or more following the offset of a 3 second light pulse (Figure 3g), a well established characteristic of the melanopsin photoresponse^1,57^. Under synaptic blockade (Figure 3h-j), optotagged ipRGCs exhibited enhanced sensitivity (Figure 3i), a shorter response latency, and longer response duration (Figure 3j) when compared to other ipRGCs.

### Functional organization of ipRGCs

Intrinsic photoresponses of ipRGCs differ considerably in the dorsal retina (Figure 1), yet optotagged ipRGCs appear homogeneous in their response latency and duration when compared to non-optotagged ipRGCs (Figure 3j). To further differentiate optotagged ipRGCs from other ipRGCs, and understand the functional diversity of ipRGCs in the mouse retina we performed unsupervised cluster analysis^48,58,59^ on ipRGC intrinsic photoresponses (Figure 4). Our unbiased approach ignores morphological and molecular criteria, and classifies ipRGCs based on their light responses. We chose to restrict our analysis to the first 3 principle components that described greater than 5% of the variance (Figure 4 a,b), and this produced 8 functional clusters at the minimal Bayesian Information Criterion (BIC; Figure 4c). Under synaptic blockade, we presented 20s blue full field light stimuli at a range of light intensities between 11.5 log photons/cm^2^/s and 13.5 log photons/cm^2^/s. Clusters were then numbered in order of their light sensitivity with earlier clusters responding to lower illumination levels (Figure 4e, f).

**Figure 4:**
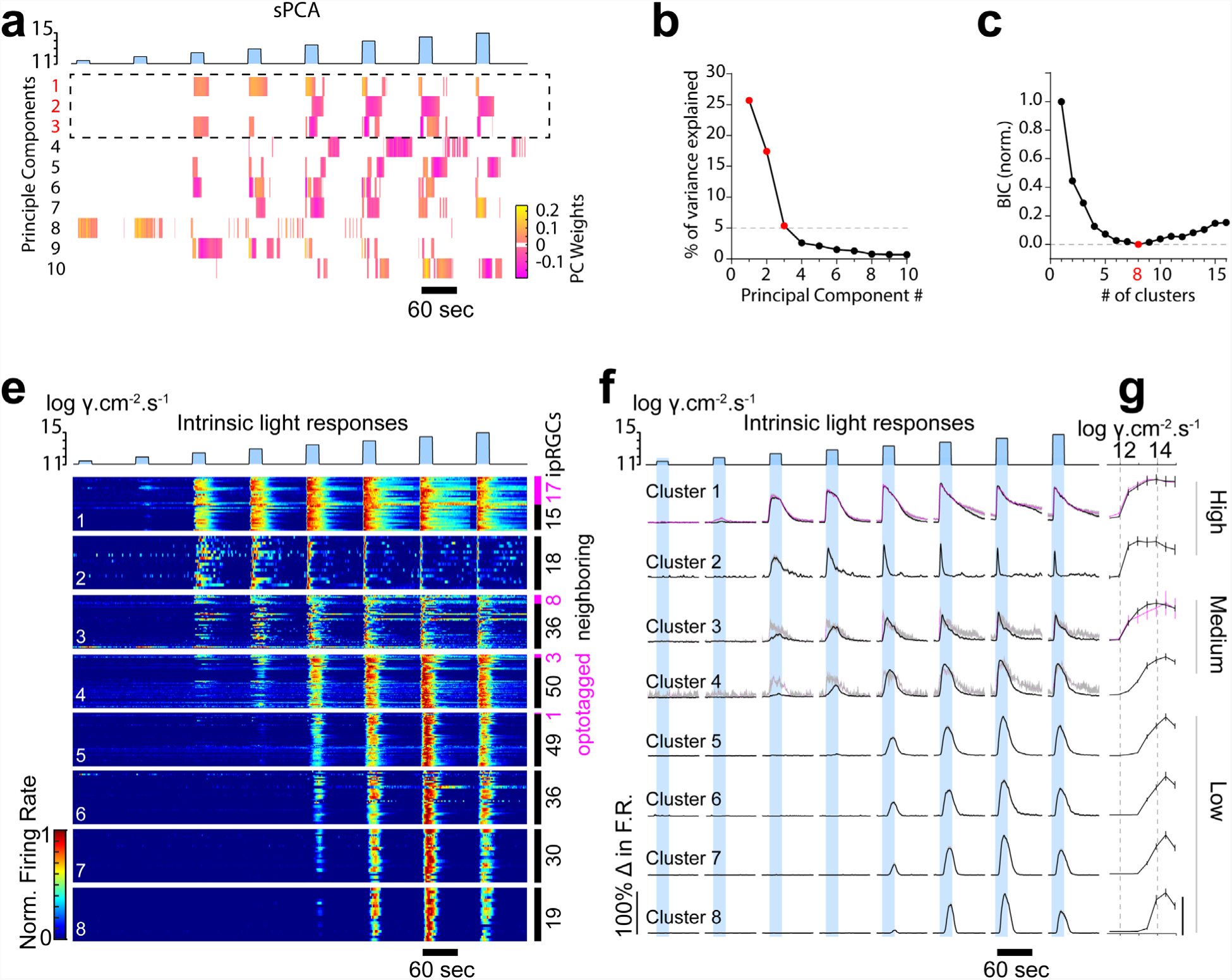
Unbiased clustering identifies eight functional clusters of ipRGCs. **(a)** Heat map of the first 10 weighted sparse principal-components generated from the intrinsic ipRGC photoresponses to increasing irradiances of 20 sec duration (blue). **(b)** Percentage of variance in ipRGC responses explained by each sparse principal component. Components that explained more than 5% of the variance were used for subsequent analysis (dotted box in a). **(c)** Normalized Bayesian Information Criterion (BIC) value plotted as a function of each potential Gaussian mixture model with a red data point (8 clusters) denoting the model chosen. BIC values serve as a balanced correlate of likelihood for the number of clusters by rewarding complexity but penalizing overfitting. The lower the BIC value, the closer the fit of the Gaussian mixture model to the data. BIC values for ipRGCs responses decline until reaching a model with 8 clustered groups (red dot), before increasing from 9 through 16. **(e)** Individual cell spike frequency heatmaps of unbiased clustered intrinsic ipRGC responses ordered by intrinsic photosensitivity. **(f)** averaged histogram of normalized intrinsic responses to increased irradiance. (e – right) Number of optotagged ipRGCs (magenta) versus surrounding ipRGCs (black) per cluster. (f – right). **(g)** Peak spike responses ± SEM for each cluster. N= 282 ipRGCs, N = 4 retina.

At a population level, ipRGCs exhibit a variable range of onsets and duration of responses to the same visual stimuli. Clusters differed in their minimum luminance sensitivity with three broad groups encompassing high, medium and low light sensitivity (Figure 4f). Cluster 1 was the most sensitive to lower light intensities followed by Cluster 2. Cluster 3 and Cluster 4 exhibited medium sensitivity with lower firing rates in response to 12.5 and 13 log photons/cm^2^/s when compared to the high-sensitivity clusters (Figure 4f). Four clusters (Cluster 5-8) were categorized as low sensitivity with three of the four clusters showing no firing to luminance between 11.5 and 13 log photons/cm^2^/s. These low sensitivity clusters qualitatively differed in the shape of their spike responses to luminance ranges between 13.5 and 15 log photons/cm^2^/s. Low sensitivity clusters had a slower rise-time and bell shaped response when compared to medium and high-sensitivity clusters, which had more rapid spiking onsets (Figure 5 a,b).

**Figure 5:**
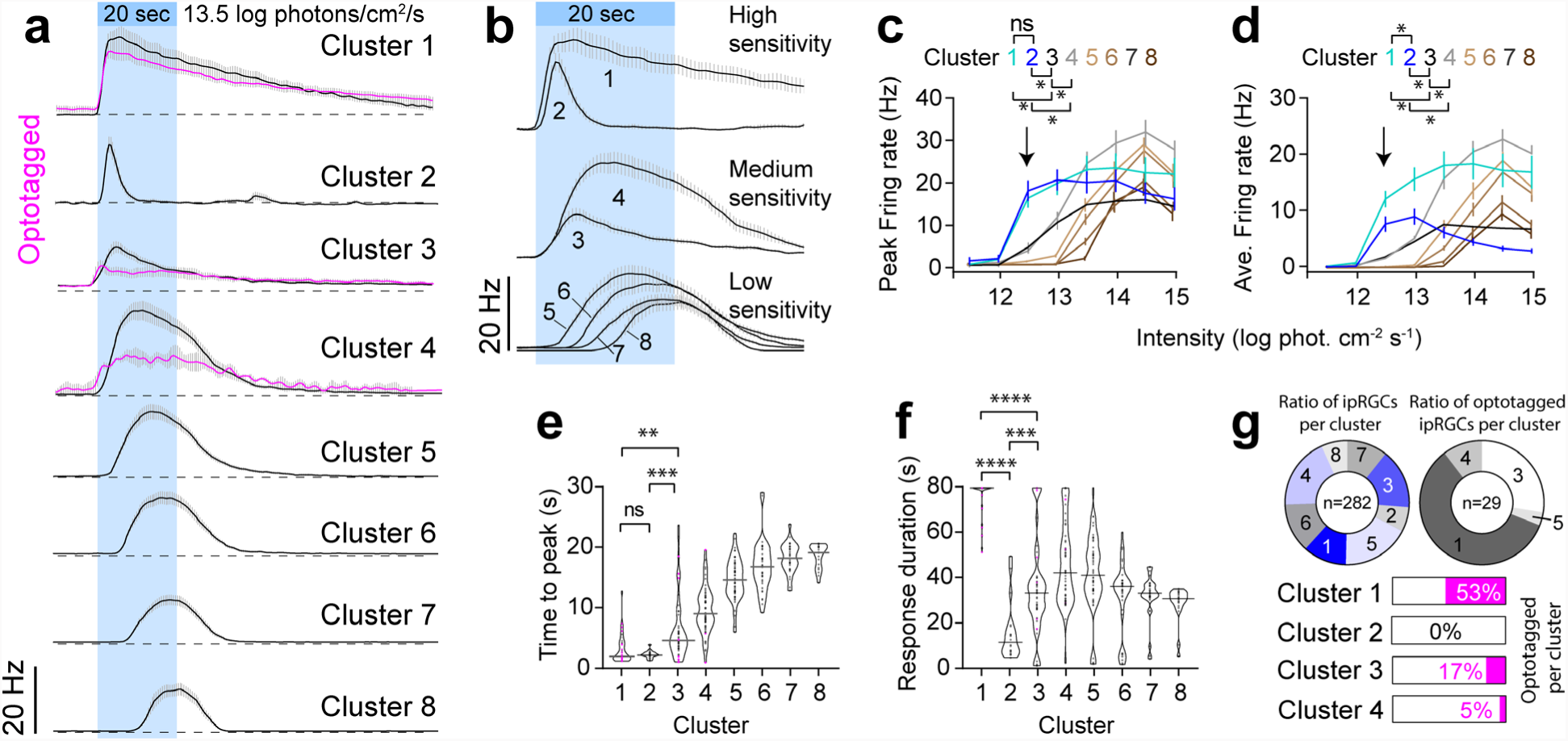
Intrinsic light responses of 8 ipRGC clusters. **(a)** Spike frequency histograms of ipRGC clusters to 20s blue illumination at 13.5 log photons/cm^2^/s. optotagged ipRGCs are plotted in magenta **(b)** Expanded traces are further grouped into high, medium and low sensitivity groups. **(c)** Peak firing rates of ipRGCs measured to blue light at increasing light intensities **(d)** Average firing rate of ipRGCs measured in response to increasing light intensities **(e)** Violin plots of time to peak for ipRGCs across all clusters **(f)** Violin plots of the response duration of spike responses of ipRGCs measured in response to 13.5 log photons cm^−2^ s^−1^. **(g)** Ratio of ipRGCs from each cluster and of ipRGCs optotaggged with ChR2 visual stimuli. Statistical significance was assessed using Mann-Whitney test for comparisons between two groups or one-way Anova with Sidaks correction for multiple comparisons (\**p*≤0.05)

We presume that the high-sensitivity clusters (Cluster 1 & 2) comprise the M1 ipRGCs, which express higher levels of melanopsin than other ipRGCs ^1,4,57,60^. The peak firing rate and the average firing rate of these clusters were highest amongst all ipRGCs to luminance of 12.5 log photons/cm^2^/s (Figure 5 c,d). As we organized these clusters by their sensitivity to illumination, their time to peak for the first level of illumination that elicited a response in all clusters (13.5 log photons/cm^2^/s) increased with cluster number (Figure 5e). The most pronounced differences however between the two highly sensitive clusters was in their response duration (Figure 5f), with cluster 1 having the longest response duration and cluster 2 having the shortest response duration, likely reflecting the impact of depolarization block on their spike responses^29^. These data provide more evidence that the highly sensitive clusters represent the M1 ipRGCs.

Of these two clusters, only one contained optotagged ipRGCs (Figure 5g), suggesting that Cluster 1 represents the SON-ipRGCs that express ChR2 using the GlyT2 promoter^9^. Two other clusters contained optotagged cells; 17% of cluster 3 ipRGCs, and 5% of Cluster 4 ipRGCs were optotagged. Both of these clusters were in the medium sensitivity group, with similar firing rates to low levels of illumination (Figure 5c, d). Given the majority of ipRGCs in the *GlyT2^Cre^;Ai32* line are display M1 or M2 morphology, we hypothesize that M2 ipRGCs are represented by one or both of these two clusters. It is also possible that a proportion of these medium-sensitive ipRGCs will be M3 ipRGCs, which are rarely encountered but have similar light responses to M2 ipRGCs^23^. The major differences between the medium sensitivity neurons is their peak firing rates and average firing rates at luminance levels of 13.5 log photons/cm^2^/s and above. Cluster 4 had the highest peak firing rate and the highest average firing rate of all ipRGCs to bright light above 14 log photons/cm^2^/s (Figure 5 c,d). This suggests that there is a possibility these two clusters represent distinct subtypes of ipRGCs with differential spike activation at depolarized membrane potentials, similar to the highly sensitive M1 ipRGCs.

Our unbiased cluster analysis identified four low sensitivity clusters that differed from the medium sensitivity clusters with a higher threshold for spike activation and a delayed time to peak (Figure 5b, d). Due to their delayed response onset and low sensitivity, we argue that these ipRGC clusters are likely comprised of M4-6 ipRGCs. Their overall spike response timing appears qualitatively similar in shape, suggesting that the primary determinant for the cluster analysis separation is due to the spike onset timing and these ipRGCs may represent similar ipRGC subtypes that have variable melanopsin expression (Figure 5 a-f). The spike response shape of the ipRGCs in these four clusters resembles previous recordings of M4 ipRGCs at similar light intensities^10,61^. It remains unclear if any of these ipRGCs represent the M5 or M6 ipRGCs, each of which expresses very low levels of melanopsin. Previous accounts of their intrinsic photosensitivity used illumination levels that are a log unit above the maximum illumination used in our recordings^24,25^ however, their overall response shape to bright blue light was similar to the lowest sensitivity clusters, suggesting that cluster 7 or cluster 8 might represent ipRGCs from M5 or M6 subtypes.

As there are at least two distinct subtypes of M1 ipRGCs in the dorsal retina that differ in their mosaic distributions^9^, we hypothesized the optotaged highly-sensitive ipRGCs fall into a subtype that differs in their intrinsic photoresponses or synaptic inputs from bipolar cells from overlapping highly-sensitive ipRGCs. However within cluster 1, optotagged ipRGCs did not have significantly different responses to their non-optotagged counterparts. This may arrive from two possible scenarios. (1) Cluster 1 includes a mixture of the same subtype containing both ChR2-expressing, and non-expressing ipRGCs. In this scenario, ChR2 expression might have been variable and detected in roughly half the SON-ipRGCs. (2) Alternatively, SON-ipRGCs and overlapping M1 ipRGCs contain the same intrinsic photoresponses and differ in other respects, such as their axon projection locations^4,13,22^, their synaptic inputs^28^, or intrinsic spiking properties^23,29^. However our data suggests there is an additional group of highly-sensitive ipRGCs, none of which fell into the optotagged cluster. With these data in mind in this second scenario, there may be three groups of M1 ipRGCs. However, it is difficult to determine the impact of blocking synaptic transmission on the spiking responses of these groups. Synaptic blockade may cause network-induced variability in the resting membrane potentials of ipRGCs and consequently impart variability in their photoresponses. Additionally, it is possible that the high and medium sensitivity groups might have differential drive from bipolar cells and that these inputs might further distinguish their light responses.

To address the effect of synaptic blockade on the clustering results, we performed retroactive analysis of the spike responses to the high and medium-sensitivity ipRGCs in response to visual stimuli presented prior to synaptic blockade (Figure 6). In these recordings we tested 535nm visual stimuli at a range of light intensities spanning scotopic to photopic illumination, to mimic bipolar cell drive mediated by both rod and cone responses (Figure 6a). Interestingly, the light-evoked spike responses of all ipRGCs were similar at light intensities up to 12 log photons/cm^2^/s, indicating their rod and cone mediated drive is similar and that their clustering is primarily driven by differences in their intrinsic photosensitivity. This model suggests that their firing rate becomes divergent at higher light intensities when melanopsin is activated or when their spike responses become blocked by depolarization as with cluster 2 (Figure 6b). These general trends occurred with green 535nm light or UV light (Figure 6C). Short 3 second blue light stimuli elicited similar responses across these groups that again diverged at higher light intensities that maximally activate melanopsin (Figure 6d-g). This divergence in responses was most apparent during sustained 1 minute blue light stimuli, illustrating the main differences between each of these clusters arises from the differential activation of melanopsin, and their spike generation mechanisms (Figure 6h-k).

**Figure 6:**
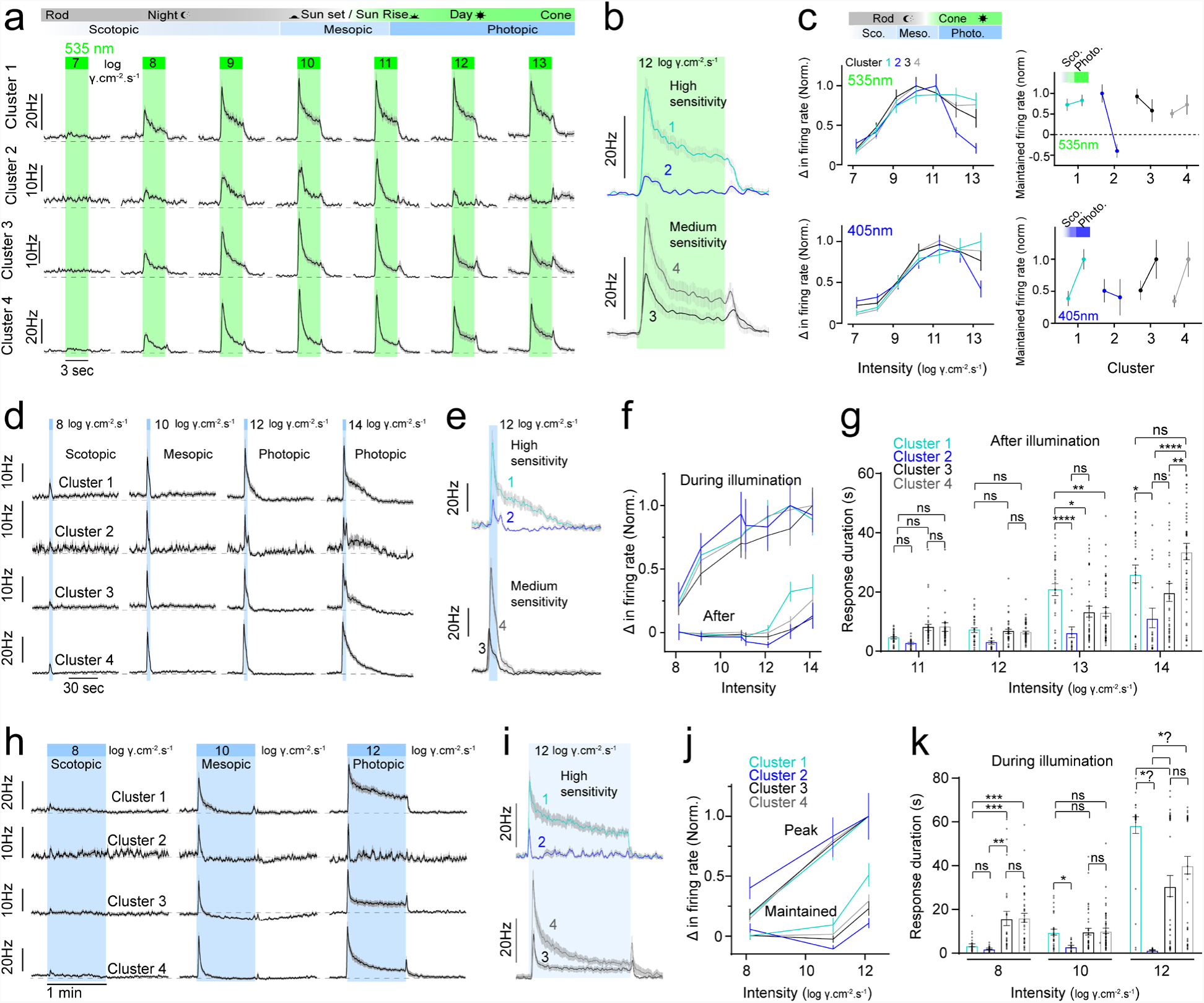
Light responses of high and medium-sensitivity ipRGCs prior to synaptic blockade. **(a)** Dark-adapted average photoresponses to green (535nm) illumination from dim moonlight (left) to bright sunlight (right) between 7 and 13 log photons/cm^2^/s. **(b)** Inset illustrating the differences in spike responses to photopic visual stimuli. Note cluster 2 ipRGCs show evidence of depolarization block. **(c)** Normalized change in firing rate across UV (blue - 405nm) and green (green - 535nm) light intensity ranges for clusters 1-4 and changes in maintained firing between scotopic and photopic illumination for each cluster (right). **(d)** Average dark-adapted photoresponses to brief blue illumination prior to synaptic blockade (3 sec). **(e)** Inset illustrating spike responses of high and medium sensitivity clusters to photopic 3s illumination. (f) Normalized change in firing rate during (top) and after (bottom) 3s blue illumination from scotopic to photopic illumination. **(g)** Bar graphs of response duration following brief illumination under mesopic to photopic light intensities. **(h)** Average dark-adapted photoresponse to 1min illumination at scotopic, mesopic and photopic light intensities. **(i)** Normalized peak (top) and maintained (bottom) change in firing rate in response to 1min visual stimuli. **(k)** Bar graphs of response duration during extended illumination from scotopic to photopic light intensities. Values are mean ± SEM. Statistical significance was assessed using Mann-Whitney test for comparisons between two groups or one-way Anova with Sidaks correction for multiple comparisons (\*\**p*≤0.01, \*\*\**p*≤0.001, \*\*\*\**p*≤0.0001).

To further explore the photoreceptor mediated inputs to ipRGCs, we analyzed the light responses to light stimuli that varied in temporal frequency and contrast at a range of light intensities^50^ (Figure 7). Because we had full datasets for all ipRGCs to both their intrinsic photoresponses and photoreceptor-mediated responses (Figure 7 e-g) we asked if performing unbiased cluster analysis using both intrinsic light responses and photoreceptor-mediated inputs would provide more resolution to uniquely cluster ipRGCs into distinct subtypes. If unique subtypes of ipRGCs differ significantly in their photoreceptor-mediated inputs, we hypothesized that this analysis would uncover additional clusters. In addition, we hypothesized that these visual stimuli might also uncover shared photoreceptor-mediated drive between ipRGC subtypes.

**Figure 7.**
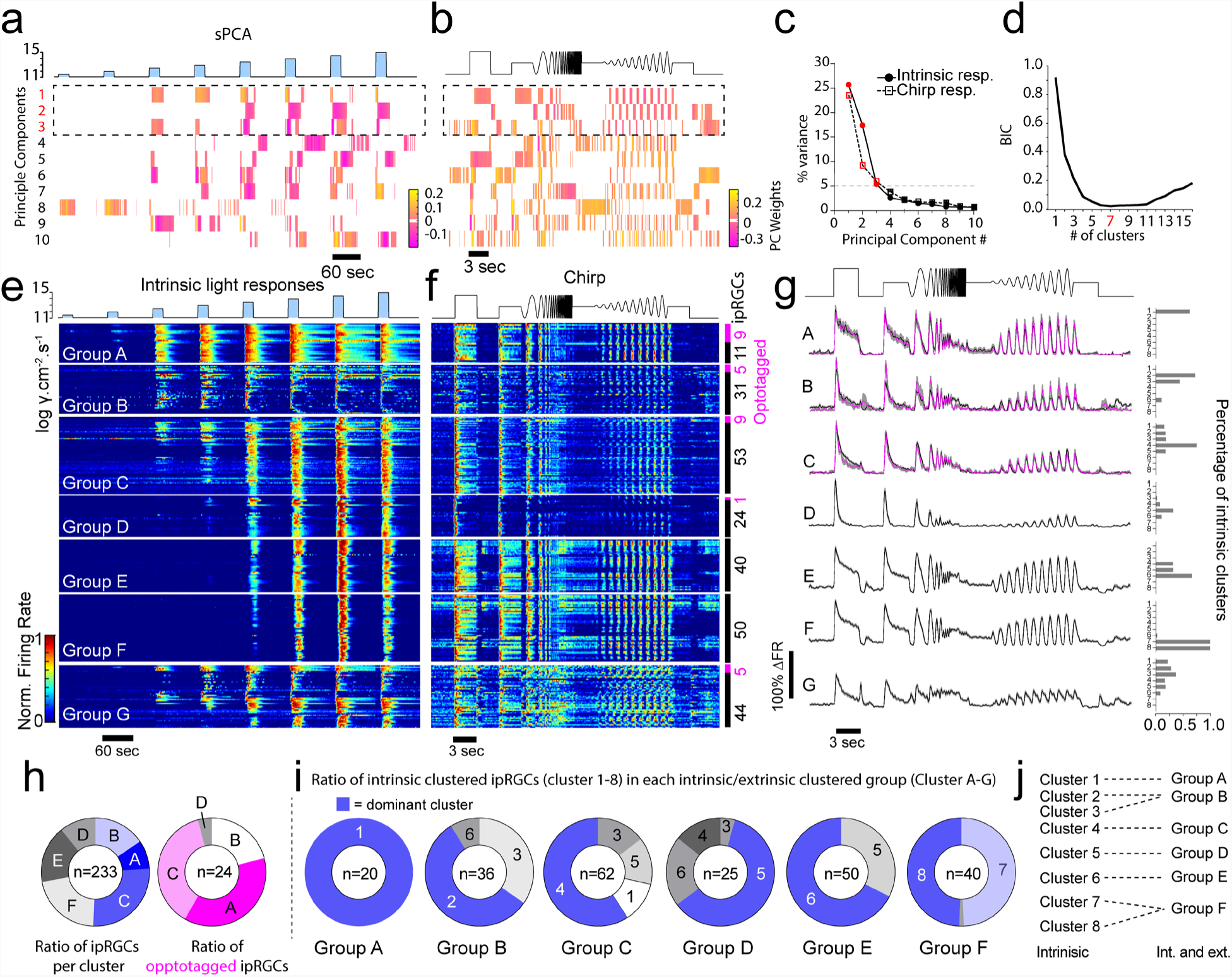
Unbiased cluster analysis of ipRGCs based on both intrinsic photoresponses and photoreceptor-mediated responses. **(a-d)** We used the first 3 principle components that explained greater than 5% of the varience resulting in the minimal BIC number of 7 clusters (d). **(e)** Spike-frequency plots of the intrinsic photoresponses and **(f)** photoreceptor-mediated responses to chirp stimuli from the 7 ipRGC groups ordered according to their intrinsic photosensitivity. optotagged ipRGCs are reporesented in magenta. **(g)** Spike frequency histograms of the 7 ipRGC groups in response to chirp stimuli. optotagged responses are plotted in magenta. The right panel represents the percentage of intrinsic clusters represented in each group. Group G represented ipRGCs that were poorly sorted and were excluded from further analysis. **(h)** Ratio of the number of ipRGCs in each group and the number of optotagged ipRGCs in each group. **(i)** Ratio of intrinsic clusters that dominate each ipRGC group. **(j)** Primary intrinsic ipRGC clusters that are represented in each of the ipRGC groups.

For clarity, we refer to these additional clusters as groups (Group A-F) to distinguish them from the intrinsic clusters (Cluster 1-8). Our results suggest that there are fewer groups of ipRGCs when clustered using these extended data (Figure 7a-g) with the minimal BIC number (Figure 7d) arising at 7 unique groups. Group G contained ipRGCs that were poorly sorted due to ambiguous spike sorting of their photoreceptor-mediated responses (Figure 7 e-g) and were removed from further analysis. We next asked how each of the ipRGC clusters arising from their intrinsic photosensitivity were arranged in each of these subsequent groups (Figure 7 g-j). Like the intrinsic ipRGC clusters, we organized each of the groups according to their intrinsic photosensitivity. The majority of Group A ipRGCs were made up of ipRGCs in the highest sensitivity cluster (Cluster 1). The dominant intrinsic cluster represented in each of the groups followed a similar pattern with the highly sensitive ipRGCs represented predominantly in Group A and B, the medium sensitive clusters represented in Group B and C and the low sensitive clusters represented in Group D, E and F.

While the assignment of ipRGC groups appears dominated by their intrinsic photosensitivity, their arrangement appears to point to shared photoreceptor-mediated drive. Medium-sensitivity ipRGCs (Cluster 3 and 4) and low sensitivity ipRGCs (Cluster 5-8) are prominent in Group C and D suggesting ipRGCs in these groups share common photoreceptor-mediated synaptic drive. Group E and F are primarily comprised of low sensitivity clusters, suggesting their clustering weights were driven primarily by their intrinsic photosensitivity however, these ipRGCs have robust responses to photoreceptor-mediated light responses at low light intensities (scotopic and mesopic) both to temporal and contrast-mediated modulation.

We next asked in these new clustered groups, how do their light-evoked spike responses driven to activate temporal frequency and contrast modulation change across background light intensities? This question is important as some ipRGCs such as the M4 ipRGCs encode changes in contrast and their melanopsin activation enhances this contrast modulation^8,10^. In addition, different M1 ipRGCs receive variable rod-mediated photoreceptor drive and melanopsin activation^28^. We examined the temporal frequency and contrast modulation of ipRGCs at different background light intensities (Figure 8). Each group had unique photoreceptor-mediated light responses to photopic, mesopic, and photopic illumination. Of the groups with the highest intrinsic photosensitivity Group B had the weakest photoreceptor-mediated responses to all light intensities (Figure 8 a,c-f). The spike responses of Group B were lower across all parameters of the stimulus space. This can be explained as Group B contained all of Cluster 2 ipRGCs (Figure 7i), which have the lowest spike frequencies and undergo depolarization block in response to bright light. Their spike responses to photoreceptor-mediated drive appear to similarly drive depolarization block.

**Figure 8.**
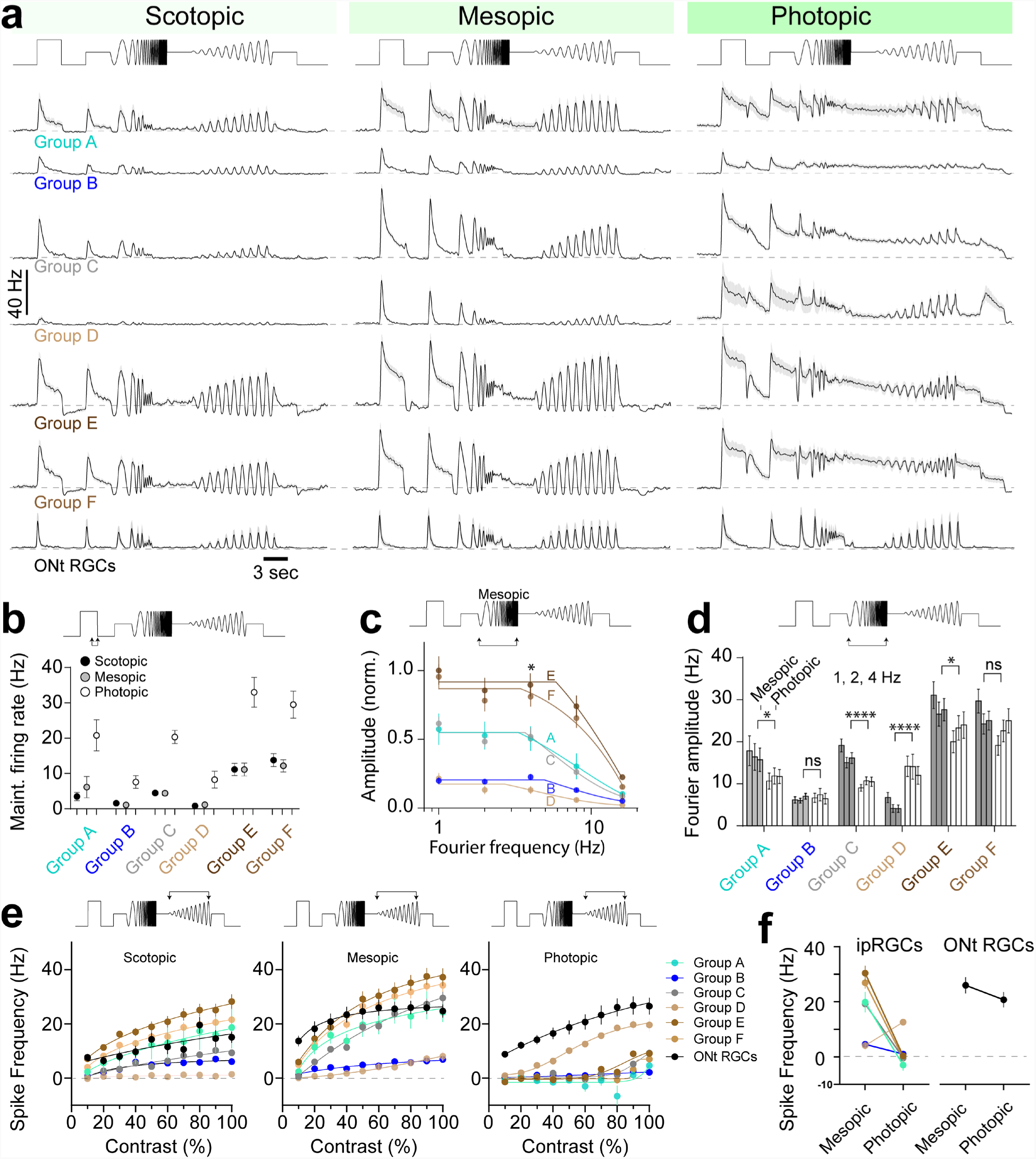
Diverse photoreceptor-mediated responses of ipRGC Groups which poorly encode visual stimuli at photopic light intensities. **(a)** Photoreceptor-mediated responses of ipRGC groups to chirp stimuli at scotopic, mesopic, and photopic light intensities. (b) Maintained firing rate taken at 3s following the onset of the first step in the chirp stimuli from ipRGCs at each light intensity. (c) Normalized fourier frequency amplitude taken from light responses to the frequency chirp component of the visual stimulus at mesopic light intensity. Group E and F had the highest frequency encoding followed by A and C. Group B and D contained the poorest frequency encoding. (d) Fourier amplitudes taken from spike responses to the frequency component of the chirp illustrate that most ipRGCs illustrate decreased frequency encoding between mesopic and photopic illumination. Group D however showed increased frequency encoding at photopic light intensities. (e) Spike frequency measurements taken from the peak responses to each contrast increment in the contrast component of the chirp. Most ipRGC groups show an increase in firing rate to increasing contrast at scotopic and mesopic light levels. Only one ipRGC group (Group D) and non-ipRGC ON transient RGCs showed robust contrast encoding at photopic light levels. (f) Spike frequency measurements between mesopic and photopic light levels (60 % contrast), measured from just prior to the contrast chirp, illustrate most ipRGCs show a marked reduction in spike rate at photopic light intensities whereas ON transient RGCs and Group D ipRGCs do not.

Group D was dominated by low sensitivity ipRGCs and had weak responses to scotopic and mesopic stimuli (Figure 8 a, e,f). The dominant ipRGCs within Group D belonged to Cluster 5 and 6. Interestingly Group E, which had robust visual responses at scotopic and mesopic light intensities is represented entirely by Cluster 5 and 6. This suggests that Group D and E represent separate populations of ipRGCs that share similarities in their intrinsic photosensitivity and melanopsin transduction, but differ substantially in their synaptic inputs.

Surprisingly, almost all ipRGC Groups were optimally activated by temporal frequency modulation and contrast modulation at mesopic light intensities (Figure 8a; Supplemental Figure 2). This is likely because melanopsin activation maintained spiking (Figure 8b) that outlasted the faster temporal components of the stimuli mediated by photoreceptors. ON transient RGCs however, which are not intrinsically photosensitive, had robust responses to photopic stimuli, similar to their scotopic and mesopic responses (Figure 8a), indicating that decreased temporal frequency modulation of ipRGCs did not arise from photoreceptor desensitization.

We measured the ability of ipRGCs to follow temporal frequency modulation using Fourier analysis of their spike responses to the chirp component of the visual stimulus (Figure 8c). The two groups with the most robust temporal frequency modulation (Group E and F) at all Fourier amplitudes contained ipRGCs with the lowest intrinsic photosensitivity. This aligns with the idea that ipRGCs expressing low levels of melanopsin are driven strongly by their synaptic inputs, reflecting their image-forming visual functions. Likewise, we might expect that the groups with the highest intrinsic photosensitivity poorly encode higher temporal frequencies. However, at meisopic light intensities, the most sensitive ipRGCs (Group A) exhibited stronger synaptic modulation than other less sensitive ipRGCs (Group D). These results are consistent with M1 SON-ipRGCs receiving strong rod-mediated drive^28^ and that some ipRGCs thought to underlie image-forming tasks (M4-6) receive little synaptic drive in low light. By measuring the Fourier frequency amplitudes of the chirp response at different light levels, we demonstrate that almost all ipRGCs significantly decrease their frequency encoding between meisopic and photopic light intensities (Figure 8d). This was with three exceptions; Group B ipRGCs, which poorly encode temporal frequencies at all light levels, remain unchanged. Group D ipRGCs, which also poorly encode temporal frequency modulation at low light levels, increased their encoding at photopic light levels. Group F ipRGCs showed a similar trend of decreased encoding but it was not significant (Figure 8d).

We next measured the ability of ipRGC groups to encode changes in contrast by measuring the peak spike frequency evoked at each contrast level in a 1Hz sinusoidal visual stimuli (Figure 8 e). These responses were measured across three light levels from scotopic to photopic. Most ipRGCs increased their spike output to increases in contrast at low light levels (scotopic and mesopic) however only Group D ipRGCs and ON transient RGCs responded to contrast modulation at photopic intensities (Figure 8 e,f). This illustrates that the maintained firing of most ipRGCs blocks their ability to encode photoreceptor-mediated contrast changes. This is surprising, particularly for M4 ipRGCs as melanopsin was shown to directly enhance their contrast responses^8,10^. This can be explained in part by differences in the experimental conditions with previous studies using visual stimuli optimized to engauge the excitatory center of these cells and our visual stimuli are full field. We did notice some spike modulation of low sensitivity ipRGCs that resemble M4 ipRGCs indicating transient decreases in spiking at photopic light intensities (Figure 8a). These occurred both in step responses at the start of the chirp and in negative contrast changes during frequency and contrast chirps. The recovery from these pauses in spiking often resembled OFF responses however, when more closely examining these responses, it appears that they arise from transient crossover inhibition (Supplementary Figure 3). These results suggest that the mechanisms of spike frequency modulation of many ipRGCs at photopic light intensities when melanopsin activates sustained firing^62^, including contrast modulation^8,10^, might not arise due to enhanced contrast encoding from melanopsin but from inhibitory modulation of their light-evoked spiking.

Together our results indicate that most ipRGCs strongly encode photoreceptor-mediated visual stimuli at low light intensities but switch to encoding light intensity at irradiances that maximally activate melanopsin.

## Discussion

We provide a comprehensive analysis of the functional diversity of ipRGCs in the mouse retina, illustrating that ipRGCs fall into different functional groups depending on the context of their visual stimuli. A key finding in the study is that the functional grouping of ipRGCs varies drastically depending on whether the cells are responding to light via inputs from their photoreceptors activated by patterned visual stimuli as compared to their grouping when activated by melanopsin. ipRGCs encode distinct temporal bandwidths driven by upstream photoreceptors in a context dependent manner, contingent upon absolute light level, driven by melanopsin and their intrinsic spiking mechanisms. Our results suggest ipRGCs respond well to image-forming visual stimuli at higher temporal frequencies and encode contrast changes at low light levels. Their bandwidth shifts towards encoding non-image forming vision, encoding or reporting overall luminance at lower temporal frequencies during daylight. Together these results illustrate an unexpected diversity in the multiplexing of visual stimuli by ipRGCs as light-intensity changes.

ipRGCs fell into three major groups (high, medium and low sensitivity) based on their intrinsic photosensitivity. We postulate that the high-sensitivity ipRGCs are M1 ipRGCs as they shared M1-like features^63^, which include high intrinsic photosensitivity (half max response ~ 12.2 log photons cm^−2^ s^−1^) and short response latency (~ 2 sec), both signs of high melanopsin expression^1,4,57,60,64^. Two of these high-sensitivity clusters differed significantly in their response duration, with Cluster 1 having the most sensitive and sustained photoresponses of all ipRGCs. Our optotagging results suggest that these ipRGCs are SON-ipRGCs that form the primary retinal projection to the supraoptic nucleus^9^. Cluster 2 had similarly sensitive responses and fast spiking onset times but only responded transiently to most visual stimuli that activated melanopsin in conditions where synaptic transmission was either intact or blocked. This transient firing and complete cessation of firing at high levels of illuminations resemble depolarization block exhibited by some ipRGCs^29^. Our results also suggest the division into unimodal and monotonic groups^29^ is present when synaptic transmission remains intact. It also demonstrates that M1 ipRGCs fall into discrete subtypes that encode the onset and persistence of illumination and that these subtypes likely project to unique sub-regions of the SCN and hypothalamus. Our results indicate that SON-ipRGCs, which are present only in the dorsal retina are monotonic, increase their spike rates to all levels of illumination. Unimodal ipRGCs, which enter depolarization block were more numerous in the ventral retina in a previous study^65^, however Cluster 2 ipRGCs were less frequently encountered in our recordings when compared to Cluster 1 ipRGCs (47 ipRGCs Cluster 1 vs 18 ipRGCs Cluster 2). This could be due to the difficulty in detecting these ipRGCs, which have smaller amplitude action potentials and lower firing rates, possibly representing lower availability or density of sodium channels. Unimodal ipRGCs may therefore be underrepresented in our MEA recordings which rely on the detection of extracellular action potentials. Alternatively, unimodal M1 ipRGCs may be more numerous in the ventral retina compared to the dorsal retina.

The remaining clusters of ipRGCs falling into medium and low sensitivity groups are likely non-M1 ipRGCs, with the medium sensitivity groups having similar properties to M2 and M3 ipRGCs. M2 ipRGCs express lower concentrations of melanopsin than M1 ipRGCs^4,60^ and M3 ipRGCs appear to have similar lower melanopsin expression and photoresponses to M2 ipRGCs^19,23^. M2 ipRGCs have tenfold lower photosensitivity than M1 ipRGCs^57^, and both M2 and M3 ipRGCs have a fast onset and sustained firing in the presence of synaptic blockers^23^. Our recordings however, cannot distinguish the two morphological types and both M2 and M3 ipRGCs may cluster similarly based on their intrinsic photosensitivity. More of our optotagged ipRGCs, which presumably contain more M2 ipRGCs^9^ fell into Cluster 3 however, some of the spike response properties of Cluster 3 M3 ipRGCs described previously^23^. Cluster 3 is more transient and has lower spiking rates at higher illumination than Cluster 4, resembling the intermittent spike frequency dependent depolarization block previously observed for M3 ipRGCs^23^. It is also possible that Cluster 3 and Cluster 4 represent different subtypes of M2 ipRGCs intermixed with M3 ipRGCs. Indeed, our Neurobiotin fills in *GlyT2^Cre^;Ai9* mouse retina illustrate M2 ipRGCs are similar to SON-ipRGCs and form territorial mosaics (Figure 2), indicating there may be multiple subtypes of M2 ipRGCs as with M1 ipRGCs.

There were 4 clusters of ipRGCs that fell into the low sensitivity groups which we presume encompass M4-6 ipRGCs which have very low levels of detectable melanopsin immunoreactivity, requiring higher light intensity levels at more extended durations to initiate melanopsin-mediated photoresponses^19,24,63,64^. Our unbiased clustering approach likely split these ipRGCs into 4 groups due to their high variability in intrinsic photosensitivity, despite having similarly shaped photoresponses (Figure 7). This bell-shaped response to blue light in the presence of synaptic blockers is similar to responses recorded previously in M4 ipRGCs^10^ however, there are few recordings of intrinsic photocurrents from M5 and M6 ipRGCs to lower levels of illumination^24,25^. It remains unclear from our analysis of spike responses under synaptic blockade if M5 and M6 ipRGCs are represented in our recordings, however they may encompass the lowest photosensitivity groups in Cluster 7 and 8.

We extended our analysis to further examine the spike responses of all ipRGCs prior to synaptic blockade by examining spiking responses of the same units on each electrode to a battery of visual stimuli designed to activate photoreceptor responses. The main conclusion from these recordings is as follows. First, we identified fewer clusters when we grouped ipRGCs based on their combined photoreceptor-mediated and intrinsic responses. This suggests that many ipRGCs share common synaptic inputs and have similar photoreceptor-mediated responses. This also suggests that the diversity of ipRGCs is more strongly driven by their melanopsin responses and intrinsic spiking mechanisms. Second, Group A was dominated by ipRGCs in Cluster 1, suggesting they represent a unique population of ipRGCs with different intrinsic photosensitivity and photoreceptor-mediated drive. Group A/Cluster 1’s sustained firing to step illumination prior to synaptic blockade as well as their highly sustained firing in response to bright light following synaptic blockade suggests that the most sensitive M1 ipRGCs may have more sustained bipolar cell drive, unique intrinsic spiking mechanisms or a combination of both when compared to other M1 ipRGCs.

The next two groups with ipRGCs (B and C) contained mixtures of ipRGCs in the high and medium sensitivity clusters. Group B was dominated by the second most sensitive ipRGCs (Cluster 2) but contained many ipRGCs from Cluster 3 and some from Cluster 6. Group C was dominated by ipRGCs from Cluster 4 but contained a small proportion of ipRGCs from Cluster 1, 3 and 5. Groups D, E, and F, were dominated by ipRGCs belonging to the low sensitive Cluster 5-8 with group F containing all the ipRGCs from the two lowest sensitivity intrinsic Cluster 7, and 8. This suggests that much of the clustering weight in these ipRGCs arises from their intrinsic photosensitivity but also points to components of the photoreceptor mediated drive that illustrate shared synaptic drive.

While ipRGCs from different intrinsic clusters were grouped together using chirp visual stimuli, the photoreceptor-driven light responses (Figure 8) between groups were remarkably different. For example, group D and E contained a majority of ipRGCs from the low sensitivity ipRGCs in Cluster 5 and 6. Still they had very different light responses to photoreceptor drive at all light levels. This suggests these ipRGCs belong to different subtypes that share similar intrinsic photosensitivity but have very different synaptic drive. Similarly, photoreceptor-mediated responses in Group B and C diverged, with group B illustrating the lowest firing rates and group C having robust responses at mesopic light stimuli. This suggests that ipRGCs in group B likely comprise M1 and M3 ipRGCs^23,29^, which have been shown to be prone to depolarization block when compared to M2 ipRGCs and that ipRGCs in Cluster 4 and group C likely comprise M2 ipRGCs.

Of the groups that contain the low sensitivity Clusters (D-F), Group E and Group F had photoreceptor-mediated responses resembling M4 ipRGCs, including robust responses to high-frequency stimulation, and robust contrast modulation. The functional contribution of melanopsin signaling is thought to adjust excitability and contrast sensitivity in these image-forming types^8,19^ using non-canonical signaling mechanisms^38,39^. Analysis of the spike frequency histograms of these groups at mesopic and photopic light intensities suggests that they receive strong crossover inhibition, reminiscent of M4 ipRGCs. Both of these groups were best at following high temporal frequency stimuli and low contrast stimuli at scoptopic and mesopic light intensities.

Surprisingly, most ipRGC groups poorly encoded temporal frequency modulation and contrast modulation at photopic light intensities. This appears to result directly from the activation of melanopsin at photopic light intensities. Two findings support this hypothesis; first, most ipRGC groups had sustained firing following the offset of step illumination. This firing was interrupted by a transient decline in spiking resembling crossover inhibition. Most ipRGCs maintained a high firing rate for the duration of light modulation following the second step in this visual stimulus and much of the modulation appeared to arise from similar crossover inhibitory input. Second, the light responses of non-ipRGC ON transient RGCs showed similar spike responses to scotopic and mesopic light intensities, suggesting ON bipolar cell inputs remain functional at these light intensities. This illustrates that most rodent ipRGCs signal changes in temporal frequencies and contrast at low illumination levels when they are likely foraging in nocturnal environments and that many ipRGCs switch roles to encode luminance during daylight.

Irradiance is the principal cue for synchronizing circadian regulation and gene expression^66^. This relies on photon counting during sustained levels of illumination and transitions in illumination, the latter of which might interrupt irradiance encoding. Sustained illumination levels occur during the day and night, with the most drastic of illumination transitions occurring at sunset and sunrise. Nocturnal foraging likely increases the probability that luminance encoding arises from reflected light off the ground, which is inverted by the lens onto the dorsal retina^67^. The two clusters of likely M1 ipRGCs in the dorsal retina presumably innervate the SCN, conveying different components of visual information. As SON-ipRGCs project to distinct SCN regions^9^ compared to other M1 ipRGCs, our data suggests the SCN circuitry requires at least two distinct forms of light information, which are segregated in the SCN. The SON-ipRGCs axon terminals are concentrated in the ventral and lateral portions of the SCN core region and avoid the arginine vasopressin (AVP) core. The SCN core receives a strong projection from M1 melanopsin ipRGGs.

Coupling anatomical and MEA recording data suggests a model in which portions of the SCN receive sustained input from Cluster 1/Group A ipRGCs, the central portion of the core receives transient synaptic input Cluster 2/Group B ipRGCs. These different light signals could mediate the unique expression of mPer1 and mPer2^68,69^, which is believed to mediate different phases of photic entrainment. Perhaps input to the SCN inner core, where non-SON ipRGCs project, is critical for light transitions from dark to light. This is consistent with evidence that minutes of even dim light exposure is sufficient for circadian entrainment in rodent models^70^. Alternatively, computation at the outer core of the SCN may be important for signaling sustained illumination. For example, animals exposed to different light intensities on a similar 12-hour cycle will entrain to the brighter of the two, establishing normal circadian rhythms. As encoding luminance across 7 orders of magnitude with a linear increase in spiking is inefficient, and does not occur, it is also possible that two independent populations of ipRGCs may be required to signal the entire range of luminance. In this model, the transient ipRGCs will signal a ‘mode’ of light intensity from scotopic, where they are functional, to photopic and inactivated. SON ipRGCs would then signal intensity gradients within each mode, allowing downstream areas to gather information about changes in luminance at specific times of the day.

In mice, the key zeitgeber appears at either dawn or dusk and as nocturnal animals emerge from their burrows at dusk they are exposed to brief and low light levels^71^. The highly sensitive and sustained ipRGCs in Cluster 1 and Group A likely present the most relevant light stimuli for circadian entrainment in low light levels. Because of the differences in spectral information that arise at dawn and dusk, specifically the increase in shorter wavelengths^72^, SON-ipRGCs in the dorsal retina dominated by medium-wavelength cones^9^, might allow ipRGCs to send color information to the SCN. As melanopsin is maximally sensitive to blue light, ipRGCs must encode differences in wavelength from their upstream photrereceptors. Restricting ipRGC subtypes to the dorsal or ventral mouse retina would form a substrate for such a mechanism.

Optotagged SON-ipRGCs represent a dominant input to the SON^9^, which is a neuroendocrine control region where peptidergic neurons regulate a variety of physiological functions, such as parturition, lactation, osmotic balance, and satiety^73^ through the release of oxytocin and vasopressin from their axons in the posterior pituitary, and throughout the brain. Although SON neurons receive synaptic inputs from the retina^74^, the functional role of this projection remains untested. Our results demonstrate that SON projecting ipRGCs deliver sustained environmental light information to SON neurons under most lighting conditions. This innervation might influence the functional release of peptides in the pituitary but might also influence oxytocin or vasopressin release directly. Therefore, SON-ipRGCs may underlie self-preservation mechanisms, altering satiety, urination, and lactation behaviors upon exposure to changes in luminance.

Our MEA approach provides several advantages over single-cell recordings of ipRGCs. First, melanopsin is directly activated by single, and 2 photon light excitation used to target fluorescent ipRGCs^42,75^. MEA recordings allowed us to avoid light exposure in dark-adapted retinas. Second, the slow, sustained response properties of ipRGCs^56^ are critical to their functional encoding of environmental light levels, temporal characteristics that necessitate long stable recordings. We acquired scotopic, mesopic, photopic, and intrinsic photoresponses from the same ipRGCs for recordings that lasted over 3 hours. Third, ipRGCs are sensitive to illumination history, adapting to environmental luminance ^31^, and ipRGCs have diverse physiological responses to light. This makes comparisons of photoresponses in neighboring ipRGCs using sequential single-cell targeted recordings, challenging. Our approach allowed us to simultaneously record from numerous neighboring ipRGCs responding to the same visual stimuli.

A disadvantage of the MEA technique in the retina is the inability to confirm the identity of recorded cells. To address this we optotagged ChR2-expressing ipRGCs at the end of the recording while applying synaptic blockers. Similar approaches identified ChR2-expressing neurons *in vivo*^76,77^ and the location of RGCs in MEA recordings^78^. This approach allowed physiologically relevant photoresponses to be acquired in groups of ipRGCs before exposing the retina to synaptic blockers and optotagging stimuli. We applied this paradigm to a recently described subpopulation of ipRGCs labeled in the *GlyT2^Cre^* mouse line^9^, that tile the dorsal retina, and overlap with other morphologically similar ipRGCs. Optotagging is advantageous for identifying ipRGCs due to the fast kinetics of ChR2^79,80^, allowing us to distinguish optotagged cells from other ipRGCs driven by the slower kinetics of melanopsin.

## Methods

### Multi-electrode array recordings & light stimulation

The MEA is an extracellular recording device consisting of multiple electrodes that allow action potentials of proximal neurons to be monitored and recorded simultaneously. MEA recordings in retinal explants provide a method of recording photoresponses from RGCs at a population level. The motivation for recording ipRGCs with this approach was to avoid the fluorescent light exposure used in single-cell targeting techniques, which can activate the melanopsin protein and cause light adaptation of the rod and cone photoreceptors. As MEA recordings of RGCs are non-specific, pharmacological and optogenetic approaches were added to localize the cells of interest.

*GlyT2Cre;Ai32* mice were dark adapted overnight and anesthetized with ketamine/xylazine. To help with removal of the vitreous during dissection and improve tissues proximity to the MEA electrodes, each eye received a 1µL intravitreal injection of 15 u/µL purified Hyaluronidase (Worthington Chemicals) 2.5 u/µL purified collagenase (Worthington Chemicals) in Balanced Salt Solution (BSS). Anesthetized animals were left on a heatpad for 15 mins then euthanized by cervical dislocation. Orientation of the eye was indicated by lightly marking the dorsal portion of the cornea and sclera with a felt tip marker. Eyes were removed and dissected in Ames solution (US Biological) under infrared illumination. Retinas were mounted on millicell membranes (Millipore, PICMORG50), and placed in an incubator under 95% C02 until recording (~30 min). Recordings were acquired on a 256-channel MEA-2100-System (Multichannel systems) at 20kHz using MC_Experimenter (Multichannel systems acquisition software). All experiments were conducted in the dark under dim red headlamps. Retinas were placed retinal ganglion side down on the MEA electrodes, and a MultiChannel Systems harp weight (Scientific Instruments) was placed on top. In order improve connectivity and stability of the recording, the retina was allowed to settle for 1hr under constant Ames perfusion and a gradual increase in temperature (room temp to 32°C).

Illumination was presented by (1) a custom-built LED array (Luxeon Star LEDs, Arduino controlled) with 405nm, 470nm, and 535nm LEDs, (2) a lightcrafter projector controlled by custom software Pystim (available at github: https://github.com/SivyerLab/pystim) and (3) a 483nm LED with collimator and control box (Thorlabs) for optogenetic stimulation. Stimuli were generally presented at increasing light intensities to retain photoresponses throughout the recording. See Table 2 for illumination and experimental details.

In three steps, a combination of illumination and pharmacological bath application was used to identify ipRGCs within the *GlyT2^Cre^;Ai32* retina. Step 1: illuminations were presented, and light responses were recorded from all RGCs in regular Ames media. Step 2: synaptic blockers (50µm DL-AP-5 (Tocris #0105), 40µm L-AP4 (Tocris #0103), 100µm CNQX (Tocris #0190) and 2µM ACET (Tocris #2728) were added to the Ames solution, pharmacologically isolating RGCs. Extended illuminations (20 sec) of increasing intensity were then used to identify the ipRGCs through their sluggish intrinsic photoresponses under blockade. Step 3: A melanopsin inhibitor^20,52^ (opsinamide; Sigma AA92593) was added to the synaptic blocking solution (10µm) and a high frequency flicker (18Hz) was used to identify optotagged or ChR2 expressing GlyT2-ipRGCs by their instantaneous time locked responses (Table 2).

During step 1, four different types of light stimuli were used (i-iv). (i) A full field illumination of 3 sec (470nm – Thor Labs) for 1.3×10^8^ to 1.3×10^14^ photons cm^−2^ s^−1^ at log unit steps, with 5 min of dark in between each exposure. (ii) A full field illumination of 60 sec (470nm – Thor Labs) for 1.3×10^8^ to 1.3×10^12^ photons cm^−2^ s^−1^ at 2 log unit steps. (iii) (iv) Full field illumination of 405nm then 535nm presented in 3 sec intervals from 1.7×10^7^ to 1.7×10^13^ photons cm^−2^ s^−1^ at 1 log unit steps. In step 2, extended illumination of 20 sec (470nm – Thor Labs) was used at light intensities from 7.9×10^10^ to 7.9×10^14^ photons cm^−2^ s^−1^ at half log unit steps with 90 sec of dark between exposures. During step 3 high frequency flicker (40x 5ms flashes at 18Hz over 9 repeated trials) was presented at 2×10^17^ photons cm^−2^ s^−1^ (483nm LED) (Thor Labs), the light intensity necessary for robust ChR2 activation^80^. Some ipRGCs are electrically connected to wide-field amacrine cells^81–83^, which signal over long ranges in the retina through their spiking axons. Therefore, it is possible that melanopsin-mediated responses are passed through gap-junctions, inducing spiking of some displaced amacrine cells, and leading us to falsely identify them in MEA recordings as ipRGCs. However, these types of responses are heavily attenuated, more likely in degenerative retina^43^, and are rarely encountered in other studies^84^.

### MEA data extraction and analysis

Raw voltage traces per retina were merged sequentially using Multichannel data manager (MCS) and Plexutility (Plexon), generating one file with all light responses. Voltage traces per channel were then bandpass filtered (Butterworth 100 and 3000 Hz). Spikes were extracted at a spike threshold of 4.5 SD from baseline with minor adjustments to insure appropriate extraction of spikes in channels with amplitudes at or near the deviation. Spikes were then rerecorded in Multichannel Analyzer (MCS). Channels with no connectivity or channels with small spiking amplitudes (typically below 20µV) were removed in order to reduce the size of the extraction file. Spike waveforms per channel were then template sorted into individual ‘units’ we call cells, via principal component analysis (PCA) using Plexon Offline Spike Sorter V3 (Plexon). Stringent thresholding of spikes during extraction (above), together with a large signal to noise during the recording, results in each electrode commonly identifying only 1–3 cells but with well-circumscribed localization in the PCA component space. Channels with more than one cell were individually confirmed to ensure that spikes are well attributed. Cells with ambiguous sorting or overlapping waveforms in the PCA component space were removed from the analysis. Trigger time points of visual stimuli were then synchronized using Multi-channel data manager, and cell responses were visualized in Neuroexplorer (Plexon). Spike timestamps were exported directly for graphing (Profit and Prism) or imported to MATLAB for further analysis using the Mathworks toolbox.

IpRGCs were identified within the MEA recordings through their photoresponses to physiological light intensities under synaptic blockade. In *GlyT2Cre;Ai32* mice, the ipRGC population, were further divided into GlyT2^Cre^-ipRGCs (optotagged ipRGCs, SON-ipRGCs) and surrounding ipRGCs (non-GlyT2 ipRGCs). GlyT2^Cre^-ipRGCs (13%±4.6 SEM of ipRGCs sorted) were identified as cells with fast, repeatable, and time locked responses to optogenetic stimulation (18Hz at 2×10^17^ photons cm^−2^ s^−1^) under synaptic block-positive melanopsin antagonist.

Perievent histograms of dark-adapted photoresponses and intrinsic photoresponses were generated from averaged firing rate across repeated trials. Changes in firing rate for dark-adapted photoresponses during (solid lines) and following illumination (dotted lines) were calculated by subtracting the baseline (spontaneous mean before illumination) and normalizing to the peak firing rate during illumination. Changes in firing rate for intrinsic photoresponses were calculated similarly but normalizing to the peak firing rate after the start of illumination as some ipRGCs took longer than the illumination window to reach a peak firing rate. Maintained firing rate was measured as average firing rate from 1 sec after illumination to the end of illumination. Sustained firing rate during extended illumination (60 sec) was measured as average firing rate during the entire illumination period. Time to peak was measured as time from illumination onset to peak firing rate. Response duration after illumination was measured as time from peak firing rate to return to spontaneous mean firing rate. Response duration during illumination was calculated similarly but limited to the duration of the illumination period (60 sec). Intrinsic responses following ChR2 activation (per trial) were calculated by subtracting the baseline (spontaneous mean) and normalizing to the average firing rate following 18Hz high frequency train (10 seconds). Perievent rasters for ChR2 responses display all spikes across all trials. Peristimulus time histograms (PSTH) graph the average normalized firing rate displayed as Z-score. Interspike interval (ISI) histogram was calculated as the frequency (occurrence) in spikes per second (instantaneous bin counts/(number of spike intervals in the spike train x bin size) per interspike interval across the entire optogenetic recording window (10 seconds before to 30 sec after 18Hz flicker across all trials). Interspike-interval (ISI) was shown as equivalent Hz as a comparison with stimulus frequency (18Hz). ISI mode histograms were plotted as the peak occurrence (mode) of ISI.

### Unbiased clustering of intrinsic photoresponses

The functional classification of ipRGCs using the clustering of photoresponses involves extracting sparse principle components (sPCA) then clustering these components using Gaussian mixture models. The approach is similar to previous studies that performed unsupervised clustering on calcium fluorescence transients from RGCs^58^, and recently ipRGCs of the neonatal mouse^48^. We applied similar methods to identify ipRGCs of the adult mouse retina using electrical responses recorded from the MEA.

Functional classification of ipRGCs was based on the clustering of intrinsic photoresponses to increases in full-field illumination. Each light response was trimmed to a window of 10 sec before and 80 sec after the start of illumination. All responsive cells were then combined into a single response matrix (cells x time points). Baseline firing rates were subtracted and each cell was normalized to its maximum value (cell normalization). Each column was also normalized to have a mean of zero; insuring that all changes in firing rate contribute variance equally (time point normalization). The sparse principle component algorithm (sPCA) from the SpaSM Matlab toolbox^59^ was then employed on our normalized response matrix, obtaining the first 10 sparse principle components. Visualizing the percentage of individual variance explained by each sPCA, we observed a sharp drop off of explained variance after principal component 3. Therefore we chose to restrict the subsequent steps of the analysis to the first 3 components each explained more than 5% of the variance within the response matrix. The weights of these sPCAs where then multiplied by the response matrix to obtain a matrix of weighted features (cells x sPCAs) with each sPCA representing a dimension. To cluster cells, we fit Gaussian Mixture Models (GMMs) to the feature matrix using Matlab’s fitgmdist algorithm, specifying 1-16 potential clusters. Each candidate cluster number was performed at 500 iterations (500×16 candidate clusters). To identify the optimal number of clusters, we implemented the Bayesian (Schwarz) Information Criterion (BIC), a model selection method that penalizes over-fitting of the data. We selected the GMM with 8 clusters as it represented the model with the lowest BIC. Clusters were constructed from the GMM distribution, cell identities were localized within each cluster and photoresponses across visual stimuli were extracted for further analysis.

## Author Contributions

M.H.B and B.S. designed the experiments, M.H.B performed the experiments, M.H.B., JL and B.S analyzed the data, M.H.B, JL, C.N.A, and B.S. wrote the manuscript.

## Acknowledgments

We would like to thank Greg Schwartz for critically reading the manuscript. This work was supported by EY032564, EY027202, EY032057, Lloyd Research Fund, Medical Research Fund of Oregon New Investigator grant P30 EY010572 and unrestricted departmental funding from Research to Prevent Blindness (New York, NY) to BS, NS103842 to CAN, EY031984 to MHB, and acknowledgement is made to the donors of National Glaucoma Research, a program of BrightFocus Foundation, for support of this research.

## Conflict of Interest

Authors declare no conflict of interest or competing financial interests.

---

**Supplemental figure 1.**
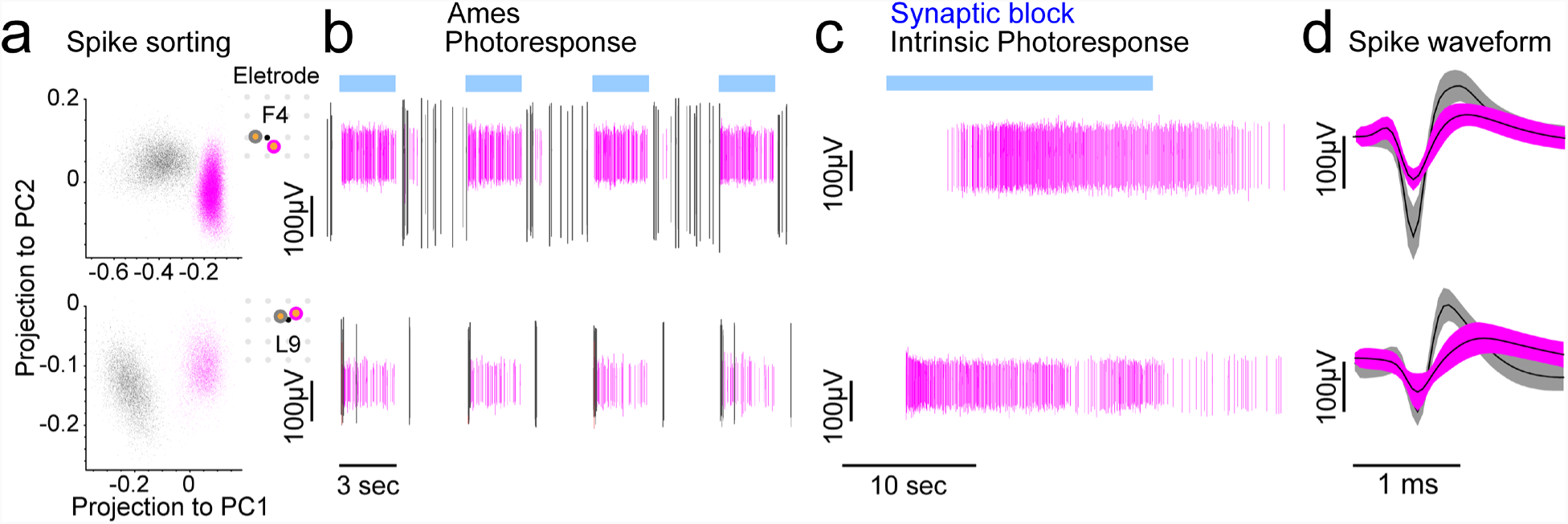
Spike sorting and example light responses of RGCs and ipRGCs. **(a)** Spike sorting is performed by using principle components analysis in Plexon Offline Spike Sorter (version 3). Two example cells from electrode F4 (top) and F9 (bottom) illustrate PCA graphs of extracted spike waveforms from a RGC (black) and an ipRGC (magenta). **(b)** Extracted spike waveforms taken in response to blue light stimuli from the cells in (a) in prior to synaptic blockade. **(c)** ipRGCs continue spiking in response to synaptic blockade. (d) Average extracted spike waveforms from the cells in (a-c).

**Supplemental figure 2.**
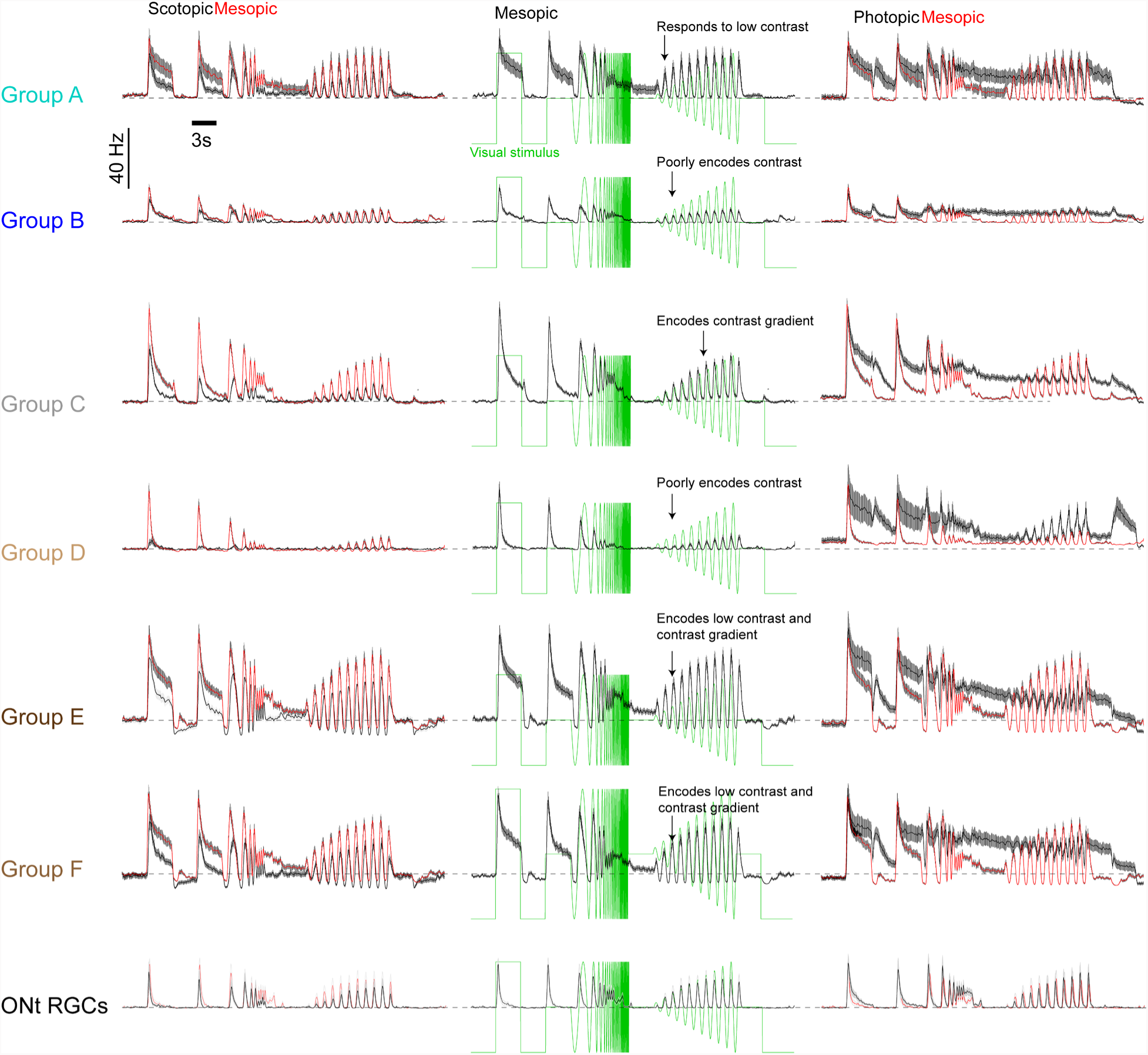
Comparisons of Scotopic, Meisopic and Photopic photoreceptor-derived light responses in ipRGC Groups and ON transient RGCs. Spike frequency histograms illustrating light responses of Group A-F ipRGCs and ON transient RGCs to photoreceptor-mediated visual stimuli at scotopic to photopic light intensities. On the left Scotopic (black) and meisopic (red) response are compared, in the middle meisopic responses are in isolation overlayed with the visual stimulus (green) and on the right photopic (black) and meisopic (red) responses are compared. Traces are averages ± SEM.

**Supplemental figure 3.**
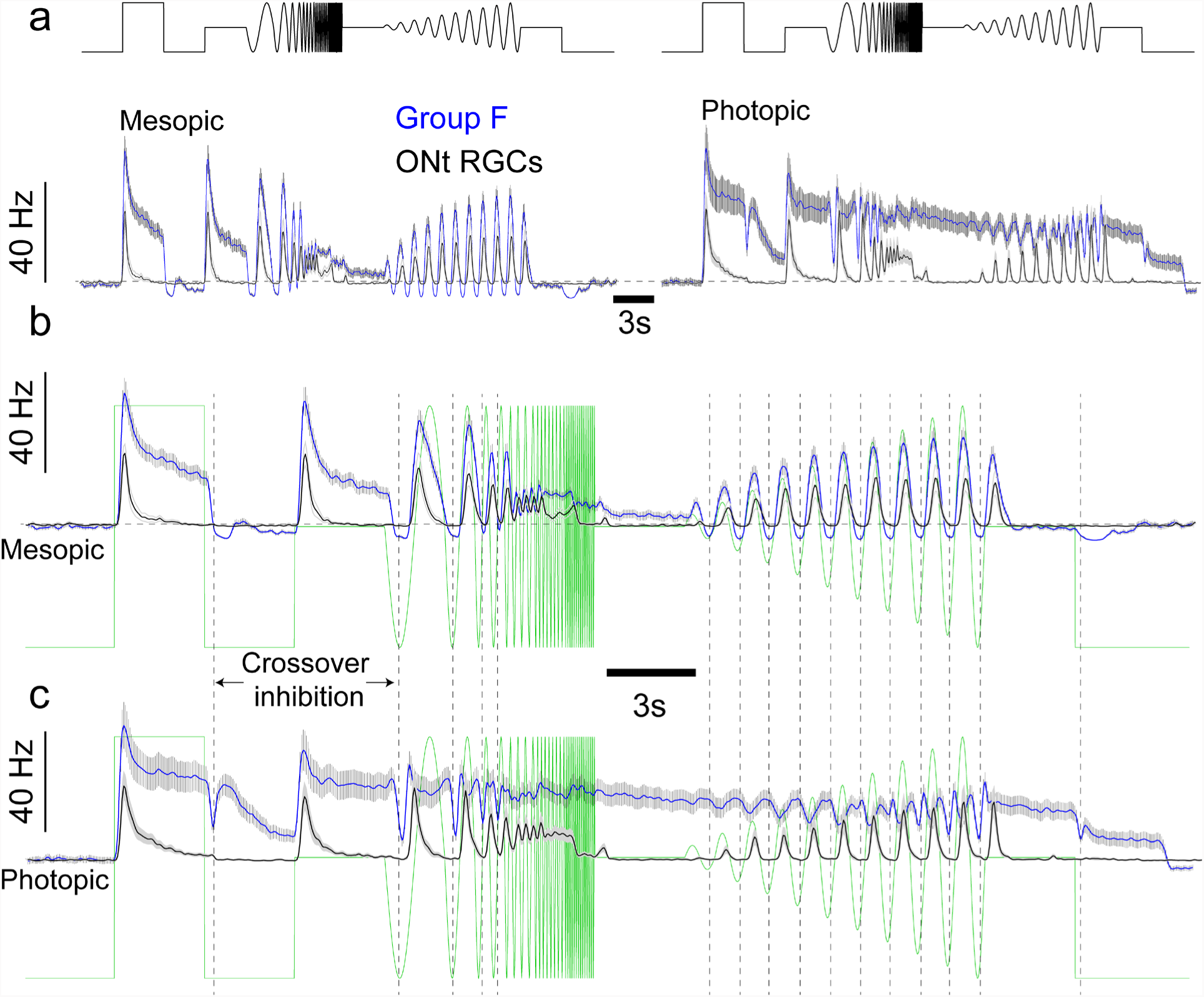
Crossover inhibition modulates spike responses of Group F ipRGCs. **(a)** Spike frequency histograms of Group F ipRGCs Blue and ON transient RGCs (black) in response to chirp stimuli at meisopic light intensities. **(b)** Expanded traces of (a) at meisopic light intensities and **(c)** photopic light intensities overlaid with the visual stimulus (green). Dashed line indicates the pause in spiking in Group F ipRGCs to negative light transitions indicative of crossover inhibition, which is particularly evident at photopic light intensities when melanopsin is maximally activated. Traces represent mean ± SEM. Crossover inhibition is not evident in ONt RGCs as their firing is transient.

## Notes

### Competing Interest Statement

The authors have declared no competing interest.

## References

1. Berson, D.M., Dunn, F.A. & Takao, M. Phototransduction by retinal ganglion cells that set the circadian clock. Science 295, 1070–1073 (2002).

2. Hattar, S., Liao, H.W., Takao, M., Berson, D.M. & Yau, K.W. Melanopsin-containing retinal ganglion cells: architecture, projections, and intrinsic photosensitivity. Science 295, 1065–1070 (2002).

3. Provencio, I., Jiang, G., De Grip, W.J., Hayes, W.P. & Rollag, M.D. Melanopsin: An opsin in melanophores, brain, and eye. Proc Natl Acad Sci U S A 95, 340–345 (1998).

4. Hattar, S., et al. Central projections of melanopsin-expressing retinal ganglion cells in the mouse. J Comp Neurol 497, 326–349 (2006).

5. Delwig, A., et al. Retinofugal Projections from Melanopsin-Expressing Retinal Ganglion Cells Revealed by Intraocular Injections of Cre-Dependent Virus. PLoS One 11, e0149501 (2016).

6. Guler, A.D., et al. Melanopsin cells are the principal conduits for rod-cone input to non-image-forming vision. Nature 453, 102–105 (2008).

7. Sondereker, K.B., Stabio, M.E. & Renna, J.M. Crosstalk: The diversity of melanopsin ganglion cell types has begun to challenge the canonical divide between image-forming and non-image-forming vision. J Comp Neurol 528, 2044–2067 (2020).

8. Sonoda, T., Lee, S.K., Birnbaumer, L. & Schmidt, T.M. Melanopsin Phototransduction Is Repurposed by ipRGC Subtypes to Shape the Function of Distinct Visual Circuits. Neuron 99, 754–767 e754 (2018).

9. Berry, M.H., et al. A melanopsin ganglion cell subtype forms a dorsal retinal mosaic projecting to the supraoptic nucleus. Nat Commun 14, 1492 (2023).

10. Schmidt, T.M., et al. A role for melanopsin in alpha retinal ganglion cells and contrast detection. Neuron 82, 781–788 (2014).

11. Lazzerini Ospri, L., Prusky, G. & Hattar, S. Mood, the Circadian System, and Melanopsin Retinal Ganglion Cells. Annu Rev Neurosci 40, 539–556 (2017).

12. Rupp, A.C., et al. Distinct ipRGC subpopulations mediate light’s acute and circadian effects on body temperature and sleep. Elife 8 (2019).

13. Chen, S.K., Badea, T.C. & Hattar, S. Photoentrainment and pupillary light reflex are mediated by distinct populations of ipRGCs. Nature 476, 92–95 (2011).

14. Fernandez, D.C., et al. Retinal innervation tunes circuits that drive nonphotic entrainment to food. Nature 581, 194–198 (2020).

15. Do, M.T.H. Melanopsin and the Intrinsically Photosensitive Retinal Ganglion Cells: Biophysics to Behavior. Neuron 104, 205–226 (2019).

16. Zhang, Z., Beier, C., Weil, T. & Hattar, S. The retinal ipRGC-preoptic circuit mediates the acute effect of light on sleep. Nat Commun 12, 5115 (2021).

17. Panda, S., et al. Melanopsin is required for non-image-forming photic responses in blind mice. Science 301, 525–527 (2003).

18. Keenan, W.T., et al. A visual circuit uses complementary mechanisms to support transient and sustained pupil constriction. Elife 5 (2016).

19. Aranda, M.L. & Schmidt, T.M. Diversity of intrinsically photosensitive retinal ganglion cells: circuits and functions. Cell Mol Life Sci 78, 889–907 (2021).

20. Mure, L.S., Vinberg, F., Hanneken, A. & Panda, S. Functional diversity of human intrinsically photosensitive retinal ganglion cells. Science 366, 1251–1255 (2019).

21. Ecker, J.L., et al. Melanopsin-expressing retinal ganglion-cell photoreceptors: cellular diversity and role in pattern vision. Neuron 67, 49–60 (2010).

22. Schmidt, T.M., Chen, S.K. & Hattar, S. Intrinsically photosensitive retinal ganglion cells: many subtypes, diverse functions. Trends Neurosci 34, 572–580 (2011).

23. Schmidt, T.M. & Kofuji, P. Structure and function of bistratified intrinsically photosensitive retinal ganglion cells in the mouse. J Comp Neurol 519, 1492–1504 (2011).

24. Quattrochi, L.E., et al. The M6 cell: A small-field bistratified photosensitive retinal ganglion cell. J Comp Neurol 527, 297–311 (2019).

25. Stabio, M.E., et al. The M5 Cell: A Color-Opponent Intrinsically Photosensitive Retinal Ganglion Cell. Neuron 97, 150–163 e154 (2018).

26. Weng, S., Estevez, M.E. & Berson, D.M. Mouse ganglion-cell photoreceptors are driven by the most sensitive rod pathway and by both types of cones. PLoS One 8, e66480 (2013).

27. Wong, K.Y., Dunn, F.A., Graham, D.M. & Berson, D.M. Synaptic influences on rat ganglion-cell photoreceptors. J Physiol 582, 279–296 (2007).

28. Lee, S.K., Sonoda, T. & Schmidt, T.M. M1 Intrinsically Photosensitive Retinal Ganglion Cells Integrate Rod and Melanopsin Inputs to Signal in Low Light. Cell Rep 29, 3349–3355 e3342 (2019).

29. Milner, E.S. & Do, M.T.H. A Population Representation of Absolute Light Intensity in the Mammalian Retina. Cell 171, 865–876 e816 (2017).

30. Emanuel, A.J., Kapur, K. & Do, M.T.H. Biophysical Variation within the M1 Type of Ganglion Cell Photoreceptor. Cell Rep 21, 1048–1062 (2017).

31. Wong, K.Y., Dunn, F.A. & Berson, D.M. Photoreceptor adaptation in intrinsically photosensitive retinal ganglion cells. Neuron 48, 1001–1010 (2005).

32. Liu, A., et al. Encoding of environmental illumination by primate melanopsin neurons. Science 379, 376–381 (2023).

33. Sonoda, T., et al. A noncanonical inhibitory circuit dampens behavioral sensitivity to light. Science 368, 527–531 (2020).

34. Johnson, E.N., et al. Distribution and diversity of intrinsically photosensitive retinal ganglion cells in tree shrew. J Comp Neurol 527, 328–344 (2019).

35. Tran, N.M., et al. Single-Cell Profiles of Retinal Ganglion Cells Differing in Resilience to Injury Reveal Neuroprotective Genes. Neuron 104, 1039–1055 e1012 (2019).

36. Berg, D.J., Kartheiser, K., Leyrer, M., Saali, A. & Berson, D.M. Transcriptomic Signatures of Postnatal and Adult Intrinsically Photosensitive Ganglion Cells. eNeuro 6 (2019).

37. Chen, L., Li, G., Jiang, Z. & Yau, K.W. Unusual phototransduction via cross-motif signaling from G(q) to adenylyl cyclase in intrinsically photosensitive retinalganglion cells. Proc Natl Acad Sci U S A 120, e2216599120 (2023).

38. Jiang, Z., Yue, W.W.S., Chen, L., Sheng, Y. & Yau, K.W. Cyclic-Nucleotide- and HCN-Channel-Mediated Phototransduction in Intrinsically Photosensitive Retinal Ganglion Cells. Cell 175, 652–664 e612 (2018).

39. Contreras, E., Nobleman, A.P., Robinson, P.R. & Schmidt, T.M. Melanopsin phototransduction: beyond canonical cascades. J Exp Biol 224 (2021).

40. Tu, D.C., et al. Physiologic diversity and development of intrinsically photosensitive retinal ganglion cells. Neuron 48, 987–999 (2005).

41. Contreras, E., Sonoda, T., Birnbaumer, L. & Schmidt, T.M. Melanopsin activates divergent phototransduction pathways in ipRGC subtypes. bioRxiv, 2022.2006.2012.495838 (2022).

42. Do, M.T., et al. Photon capture and signalling by melanopsin retinal ganglion cells. Nature 457, 281–287 (2009).

43. Eleftheriou, C.G., et al. Melanopsin Driven Light Responses Across a Large Fraction of Retinal Ganglion Cells in a Dystrophic Retina. Front Neurosci 14, 320 (2020).

44. Mouland, J.W., Watson, A.J., Martial, F.P., Lucas, R.J. & Brown, T.M. Colour and melanopsin mediated responses in the murine retina. Front Cell Neurosci 17, 1114634 (2023).

45. Sekaran, S., et al. Melanopsin-dependent photoreception provides earliest light detection in the mammalian retina. Curr Biol 15, 1099–1107 (2005).

46. Perez-Leighton, C.E., Schmidt, T.M., Abramowitz, J., Birnbaumer, L. & Kofuji, P. Intrinsic phototransduction persists in melanopsin-expressing ganglion cells lacking diacylglycerol-sensitive TRPC subunits. Eur J Neurosci 33, 856–867 (2011).

47. Kirkby, L.A. & Feller, M.B. Intrinsically photosensitive ganglion cells contribute to plasticity in retinal wave circuits. Proc Natl Acad Sci U S A 110, 12090–12095 (2013).

48. Caval-Holme, F., Zhang, Y. & Feller, M.B. Gap Junction Coupling Shapes the Encoding of Light in the Developing Retina. Curr Biol 29, 4024–4035 e4025 (2019).

49. Arroyo, D.A., Kirkby, L.A. & Feller, M.B. Retinal Waves Modulate an Intraretinal Circuit of Intrinsically Photosensitive Retinal Ganglion Cells. J Neurosci 36, 6892–6905 (2016).

50. Baden, T., et al. The functional diversity of retinal ganglion cells in the mouse. Nature 529, 345–350 (2016).

51. Keppler, A., et al. A general method for the covalent labeling of fusion proteins with small molecules in vivo. Nat Biotechnol 21, 86–89 (2003).

52. Jones, K.A., et al. Small-molecule antagonists of melanopsin-mediated phototransduction. Nat Chem Biol 9, 630–635 (2013).

53. Pant, M., Zele, A.J., Feigl, B. & Adhikari, P. Light adaptation characteristics of melanopsin. Vision Res 188, 126–138 (2021).

54. Do, M.T. & Yau, K.W. Adaptation to steady light by intrinsically photosensitive retinal ganglion cells. Proc Natl Acad Sci U S A 110, 7470–7475 (2013).

55. Mure, L.S., et al. Melanopsin-Encoded Response Properties of Intrinsically Photosensitive Retinal Ganglion Cells. Neuron 90, 1016–1027 (2016).

56. Emanuel, A.J. & Do, M.T. Melanopsin tristability for sustained and broadband phototransduction. Neuron 85, 1043–1055 (2015).

57. Schmidt, T.M. & Kofuji, P. Functional and morphological differences among intrinsically photosensitive retinal ganglion cells. J Neurosci 29, 476–482 (2009).

58. Baden, T., et al. The functional diversity of retinal ganglion cells in the mouse. Nature 529, 345–350 (2016).

59. Sjöstrand, K., Clemmensen, L.H., Larsen, R., Einarsson, G. & Ersbøll, B. SpaSM: A MATLAB Toolbox for Sparse Statistical Modeling. 2018 84, 37 (2018).

60. Baver, S.B., Pickard, G.E., Sollars, P.J. & Pickard, G.E. Two types of melanopsin retinal ganglion cell differentially innervate the hypothalamic suprachiasmatic nucleus and the olivary pretectal nucleus. Eur J Neurosci 27, 1763–1770 (2008).

61. Sonoda, T., Okabe, Y. & Schmidt, T.M. Overlapping morphological and functional properties between M4 and M5 intrinsically photosensitive retinal ganglion cells. J Comp Neurol 528, 1028–1040 (2020).

62. Mure, L.S., et al. Sustained Melanopsin Photoresponse Is Supported by Specific Roles of beta-Arrestin 1 and 2 in Deactivation and Regeneration of Photopigment. Cell Rep 25, 2497–2509 e2494 (2018).

63. Zhao, X., Stafford, B.K., Godin, A.L., King, W.M. & Wong, K.Y. Photoresponse diversity among the five types of intrinsically photosensitive retinal ganglion cells. The Journal of physiology 592, 1619–1636 (2014).

64. Reifler, A.N., et al. The rat retina has five types of ganglion-cell photoreceptors. Exp Eye Res 130, 17–28 (2015).

65. Milner, E.S. & Do, M.T.H. A Population Representation of Absolute Light Intensity in the Mammalian Retina. Cell 171, 865–876.e816 (2017).

66. Lucas, R.J., et al. Measuring and using light in the melanopsin age. Trends in neurosciences 37, 1–9 (2014).

67. Qiu, Y., et al. Natural environment statistics in the upper and lower visual field are reflected in mouse retinal specializations. Current biology : CB 31, 3233–3247 e3236 (2021).

68. Zheng, B., et al. Nonredundant roles of the mPer1 and mPer2 genes in the mammalian circadian clock. Cell 105, 683–694 (2001).

69. Shigeyoshi, Y., et al. Light-induced resetting of a mammalian circadian clock is associated with rapid induction of the mPer1 transcript. Cell 91, 1043–1053 (1997).

70. Foster, R.G., Hughes, S. & Peirson, S.N. Circadian Photoentrainment in Mice and Humans. Biology (Basel) 9, 180 (2020).

71. Foster, R.G., Hughes, S. & Peirson, S.N. Circadian Photoentrainment in Mice and Humans. Biology (Basel) 9 (2020).

72. Walmsley, L., et al. Colour as a signal for entraining the mammalian circadian clock. PLoS Biol 13, e1002127 (2015).

73. Hume, C., Sabatier, N. & Menzies, J. High-Sugar, but Not High-Fat, Food Activates Supraoptic Nucleus Neurons in the Male Rat. Endocrinology 158, 2200–2211 (2017).

74. Cui, L.-N., Jolley, C.J. & Dyball, R.E.J. Electrophysiological Evidence for Retinal Projections to the Hypothalamic Supraoptic Nucleus and its Perinuclear Zone. J Neuroendocrinol 9, 347–353 (1997).

75. Denk, W. & Detwiler, P.B. Optical recording of light-evoked calcium signals in the functionally intact retina. Proc Natl Acad Sci U S A 96, 7035–7040 (1999).

76. Madisen, L., et al. A toolbox of Cre-dependent optogenetic transgenic mice for light-induced activation and silencing. Nature neuroscience 15, 793–802 (2012).

77. Gale, S.D. & Murphy, G.J. Active Dendritic Properties and Local Inhibitory Input Enable Selectivity for Object Motion in Mouse Superior Colliculus Neurons. J Neurosci 36, 9111–9123 (2016).

78. Hilgen, G., et al. Unsupervised Spike Sorting for Large-Scale, High-Density Multielectrode Arrays. Cell Rep 18, 2521–2532 (2017).

79. Deisseroth, K. & Hegemann, P. The form and function of channelrhodopsin. Science (New York, N.Y.) 357, eaan5544 (2017).

80. Lin, J.Y. A user’s guide to channelrhodopsin variants: features, limitations and future developments. Experimental physiology 96, 19–25 (2011).

81. Harrison, K.R., Chervenak, A.P., Resnick, S.M., Reifler, A.N. & Wong, K.Y. Amacrine Cells Forming Gap Junctions With Intrinsically Photosensitive Retinal Ganglion Cells: ipRGC Types, Neuromodulator Contents, and Connexin Isoform. Invest Ophthalmol Vis Sci 62, 10 (2021).

82. Sabbah, S., Berg, D., Papendorp, C., Briggman, K.L. & Berson, D.M. A Cre Mouse Line for Probing Irradiance- and Direction-Encoding Retinal Networks. eNeuro 4 (2017).

83. Reifler, A.N., et al. All Spiking, Sustained ON Displaced Amacrine Cells Receive Gap-Junction Input from Melanopsin Ganglion Cells. Curr Biol 25, 2878 (2015).

84. Segev, R., Goodhouse, J., Puchalla, J. & Berry, M.J., 2nd. Recording spikes from a large fraction of the ganglion cells in a retinal patch. Nat Neurosci 7, 1154–1161 (2004).

